## Supplementary figures and images for "A cattle graph genome incorporating global breed diversity"

### Supplementary Figure 1

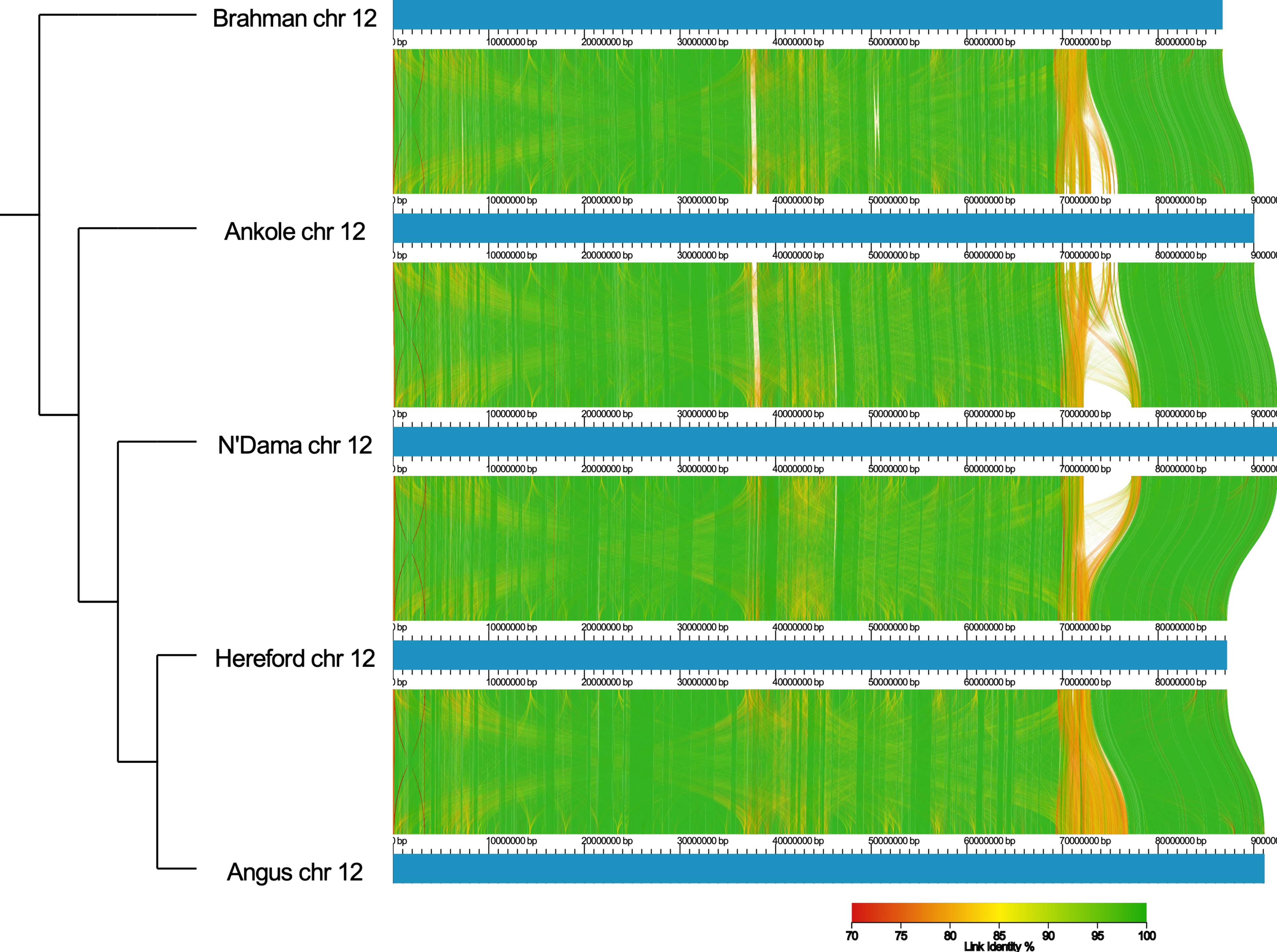

### Supplementary Figure 2

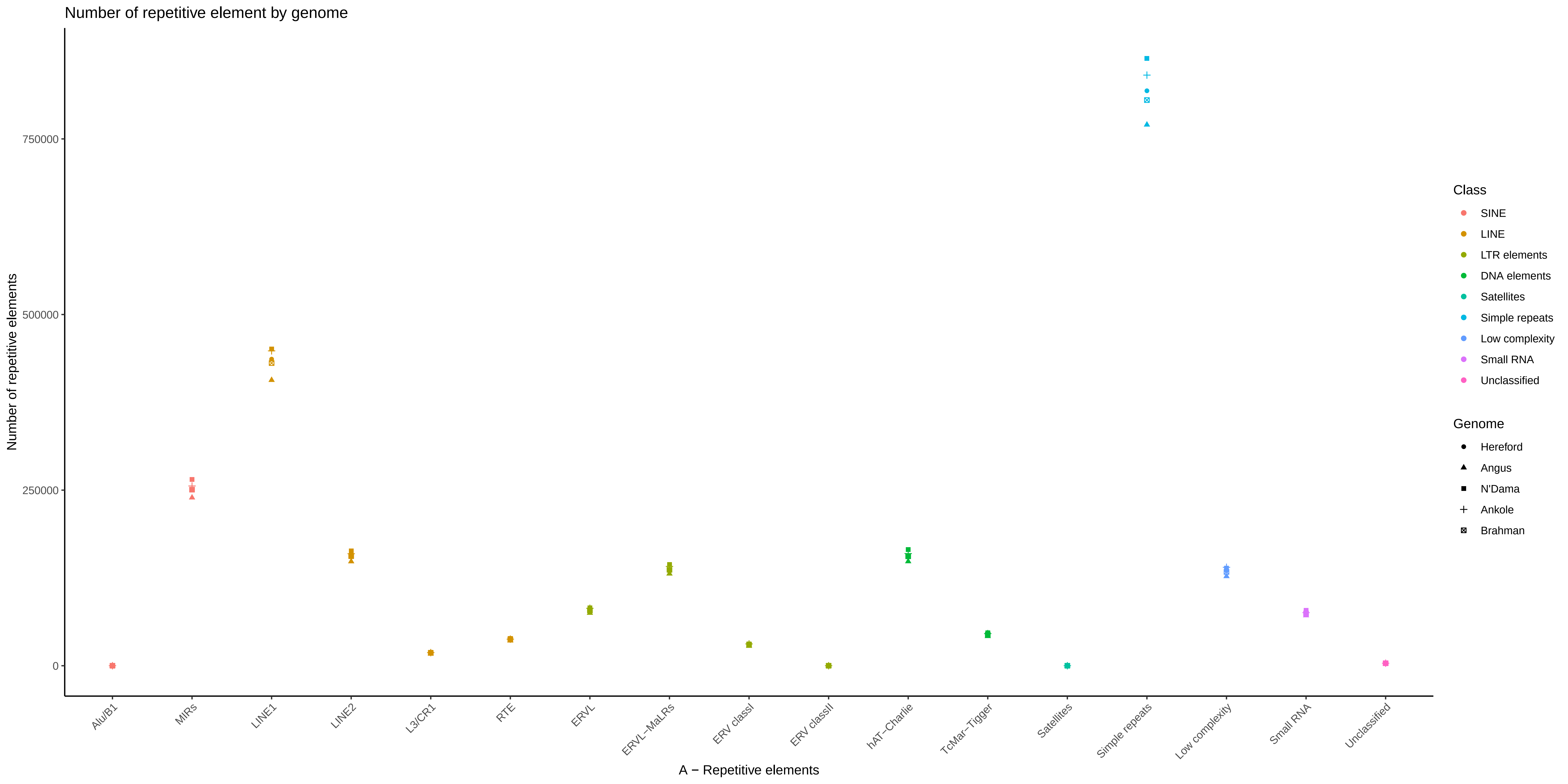

### Supplementary Figure 3

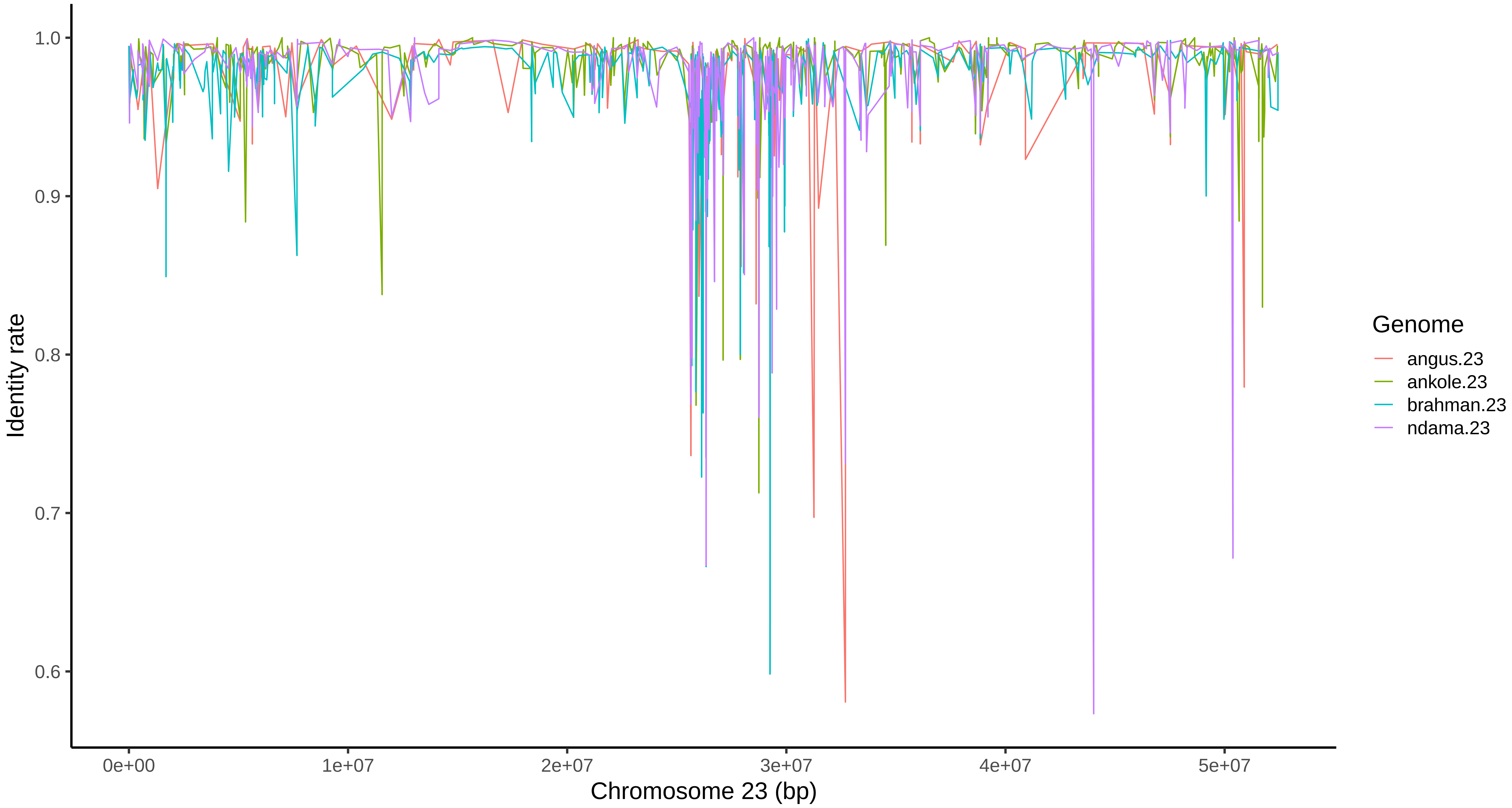

### Supplementary Figure 4

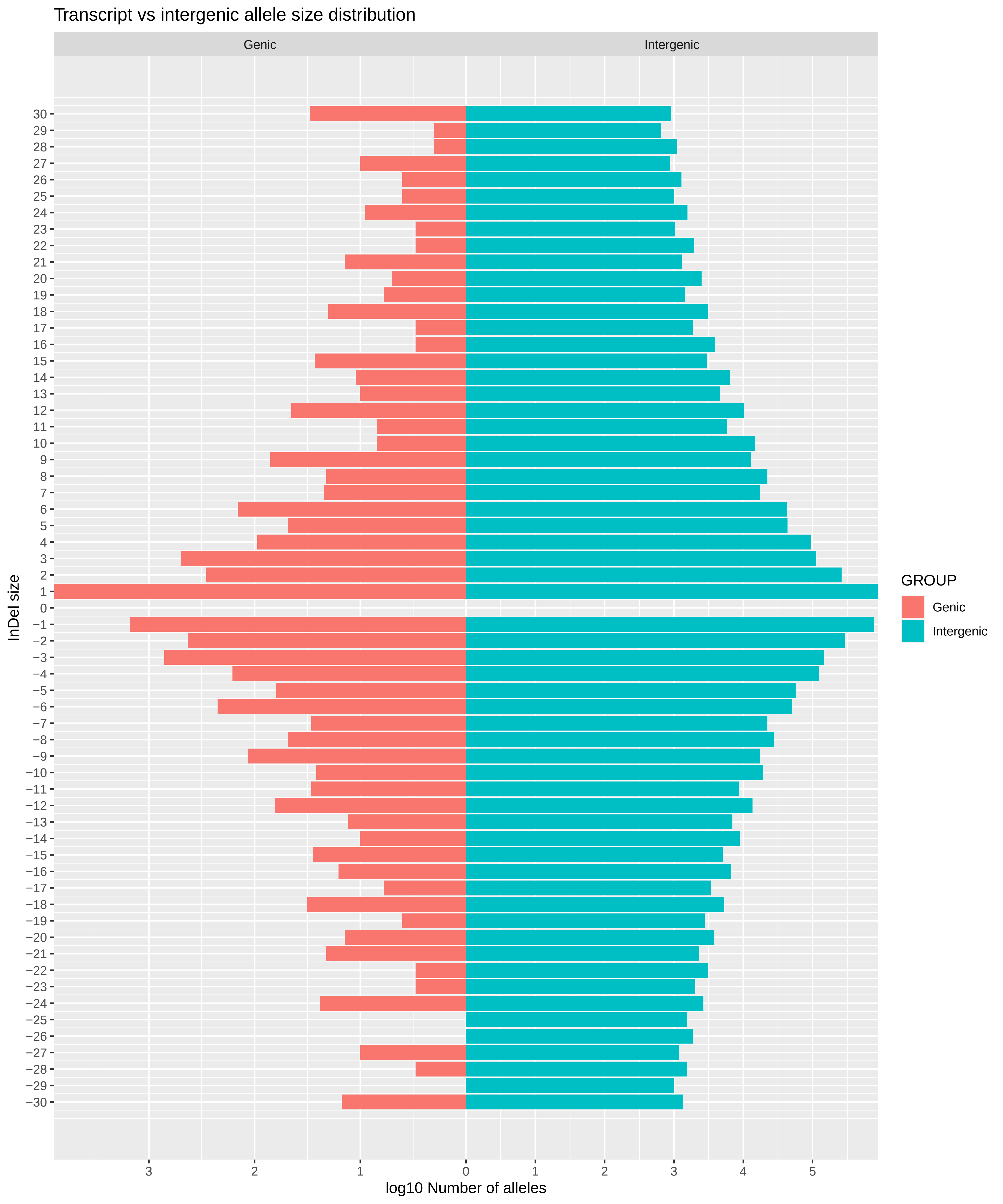
