## Supplementary Methods 1 for "A cattle graph genome incorporating global breed diversity"

**Supplementary methods 1 - ATAC-seq sample preparation**

*Extraction of PBMC DNA and isolation of B cells*

Peripheral blood mononuclear cells (PBMCs) were isolated from Holstein Friesian, N’Dama and Nelore peripheral blood by density gradient centrifugation using Ficoll Plaque Plus (GE Healthcare). DNA was extracted from PBMC using a QIAGEN DNeasy blood and tissue kit with proteinase K and RNase treatment. For isolation of B cells, PBMCs were resuspended at 3x10^7^ cells/ml in PBS/2 mM EDTA/0.5% BSA and incubated with 0.066 μg/ml ILA58 monoclonal antibody, which binds the immunoglobulin light chain, for 20 min at 4°C. After two washes in PBS/2 mM EDTA/0.5% BSA, PBMCs were incubated in 2 ml of 2 μg/ml PE-conjugated goat anti-mouse IgG2a (Molecular probes) for 20 min at 4°C. PBMCs were washed twice and were resuspended in PBS/2 mM EDTA/0.5% BSA with or without 1 µg/ml DAPI (Invitrogen). B cells were sorted on a FACSAria II or III Cell Sorter (BD Biosciences) or BD Influx Cell Sorter (BD Biosciences) with 82 - 97 % purity validated by post-sort flow cytometric analysis of 1,000 events. All cells were stained and sorted within 9 hours of blood collection and kept on ice between processing steps.

*ATAC-seq library preparation*

Cells of the mouse mastocytoma cell line P815 were spiked into the Holstein Freisian B cell sample at a 1:10 ratio. For the three breeds, 50,000 cells were transferred into a 96-well v-bottomed plate on ice. Cells were centrifuged at 500 x*g*, 4°C for 2 min and the supernatant was removed. Next, cells were resuspended in 100 µl cold lysis buffer (10 mM Tris hydrochloride, pH 7.4, 10 mM sodium chloride, 3 mM magnesium chloride, 0.1% IGEPAL CA-630). Cells were centrifuged at 500 x*g* at 4°C for 10 min and the supernatant was discarded. Nuclei were resuspended in 50 µl transposase mixture (25 μl 2x TD buffer, 2.5 μl TDE1 Tagment DNA (Illumina) and 22.5 μl nuclease-free water), transferred to 1.5 ml microcentrifuge tubes, and then were incubated for 30 min at 37°C in an Eppendorf Thermomixer with agitation at 300 rpm. Transposed DNA was purified using a QIAGEN MinElute Reaction Cleanup Kit with elution in 14 µl water. To generate presumably nucleosome-free ATAC-seq libraries, these steps were repeated using 1500 - 2000 ng Holstein Friesian PBMC DNA treated with protease K in replacement of the 50,000 cells. Transposed DNA was then amplified using Nextera primers listed in Buenrostro et al., 2015. Specifically, all samples were amplified using the index i5 primer v2_Ad1.1_TAGATCGC and one of the index i7 v2_Ad2.1 - 2.12 primers. qPCR reactions were carried out in duplicate using 0.5 µl transposed DNA, 5 µl NEBNext High-Fidelity 2x PCR Master Mix (NEB), 1.25 µl 10 µM dual-index PCR primers (Integrated DNA Technologies), 0.25 µl 20x SYBR Green I (Invitrogen), 0.15 µl 1 mM ROX reference dye (Agilent Technologies), and 1.6 µl nuclease-free water (QIAGEN). The PCR conditions were as follows: 72°C for 5 min, then 98°C for 30 sec, followed by 21 thermocycles at 98°C for 10 sec, 63°C for 30 sec and 72°C for 1 min. To calculate the appropriate number of cycles for amplification of the remaining transposed DNA, linear Rn was plotted against cycle number to determine the cycle number corresponding to one-quarter of the maximum fluorescent intensity, with an average taken across duplicates. This number of cycles was used to amplify the remaining 12.5 μl transposed DNA using 25 µl NEBNext High-Fidelity 2x PCR Master Mix (NEB), 6.25 µl 10 µM dual-index PCR primers, with the same cycling conditions as described for qPCR. The DNA was purified using a Qiagen MinElute PCR Purification Kit (QIAGEN) with elution in 20 µl nuclease-free water (QIAGEN). To remove residual primers, the libraries were purified using 1.4X AMPure beads (Beckman Coulter). Then, two further AMPure steps were performed to remove large DNA fragments (>1000bp). The first used 0.5X AMPure beads with recovery of the supernatant, to which 1.3X AMPure beads were added to purify the final libraries. Library quality was assessed for the fragment length distribution on a 2200 TapeStation instrument (Agilent Technologies), using High Sensitivity D1000 ScreenTape and Reagents (Agilent Technologies). The library concentrations were measured on a Qubit 3.0 (Invitrogen) using a Qubit dsDNA High Sensitivity assay kit (Invitrogen). Resulting libraries were sequenced using 75 bp paired-end sequencing on a HiSeq 4000 sequencer (Illumina) or 50 bp paired-end sequencing on NovaSeq 6000 sequencer (Illumina) at the Edinburgh genomics facility.

*References*

Buenrostro, J.D., Wu, B., Litzenburger, U.M., Ruff, D., Gonzales, M.L., Snyder, M.P., Chang, H.Y., and Greenleaf, W.J. (2015). Single-cell chromatin accessibility reveals principles of regulatory variation. Nature 523, 486–490.
