## Supplementary Note 1 for "A cattle graph genome incorporating global breed diversity"

Supplementary Note 1 – The N’Dama assembly report

### N’Dama long read sequencing

#### Pre-Assembly statistics of sample’s long reads

Alignment statistics to latest Bos taurus assembly via minimap2:

minimap2 -ax map-pb BtauARS.fasta Ndama.subreads.bam | samtools view - -S -h -b | samtools sort - -o ./Ndama.subreads_minimap.bam

Alignment statistics obtained through samtools flagstat.

In total, reads (87.95%) were aligned to the reference genome.

Coverage obtained on the chromosomes (samtools depth) as the average of all basis within each chromosome (Figure 1).

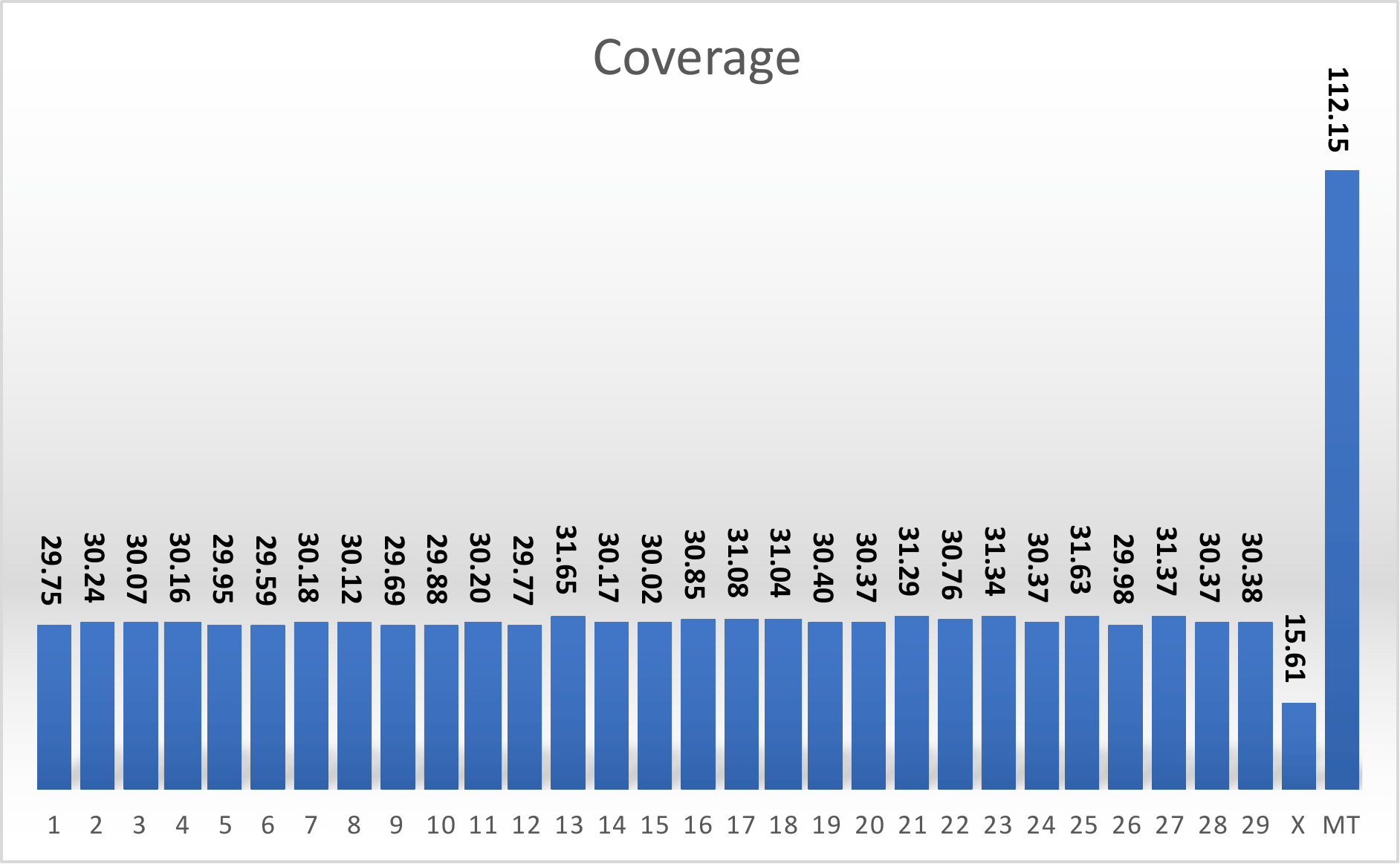

Coverage per chromosome of mapped PacBio reads

### Assembly Selection

We generated two different assemblies from two different software, CANU and FALCON. Different metrics used to select the genome for subsequent analysis.

| Parameter | CANU | FALCON |
| --- | --- | --- |
| BP yield | 2,624,985,022 | 2,644,771,833 |
| # sequences (polished) | 5,940 | 4,115 |
| N25 | 4,717,823 | 6,209,788 |
| N50 | 2,636,435 | 3,275,473 |
| N75 | 1,195,117 | 1,561,644 |
| N90 | 328,738 | 477,322 |
| N95 | 99,257 | 124,586 |
| L25 | 93 | 78 |
| L50 | 285 | 227 |
| L75 | 652 | 518 |
| Min L | 1,007 | 9,000 |
| Avg L | 442,266 | 641,623 |
| Median L | 44,046 | 69,448 |
| Max L | 21,377,347 | 18,329,644 |
| GC% | 41.95% | 42.04% |
| BUSCO Complete* | 77.90% | 85.84% |
| BUSCO Fragmented* | 14.00% | 6.99% |
| BUSCO Missing* | 8.10% | 7.16% |

Based on the metrics above, FALCON assembly have been selected since it shows a high contig metrics, bp yield and includes a filtering for highly-repetitive regions. The resulting assembly has been polished twice using long reads with Racon, and once with short reads and Pilon v1.23. Long reads were mapped using minimap2, whereas short reads were mapped using bwa mem.

Pilon polishing step allowed to fix multiple misassemblies, insertions, deletions, collapse repeated regions and trim low coverage bases. Below a short summary of the polishing step:

|  | Value |
| --- | --- |
| Original size | 2,655,094,705 |
| Bases confirmed | 2,609,970,564 |
| Bases confirmed (%) | 98.30% |
| Corrected size | 2,644,876,474 |
| SNPs corrected | 593,486 |
| Insertions corrected | 5,129,742 |
| Insertions corrected (bp) | 8,254,027 |
| Deletions corrected | 566,769 |
| Deletions corrected (bp) | 686,240 |
| Collapsed bases | 11,721,029 |

After running Pilon on the assembly created using Falcon-Unzip, we run a completeness assessment using BUSCO v3, with the assembly showing a good completeness (91%).

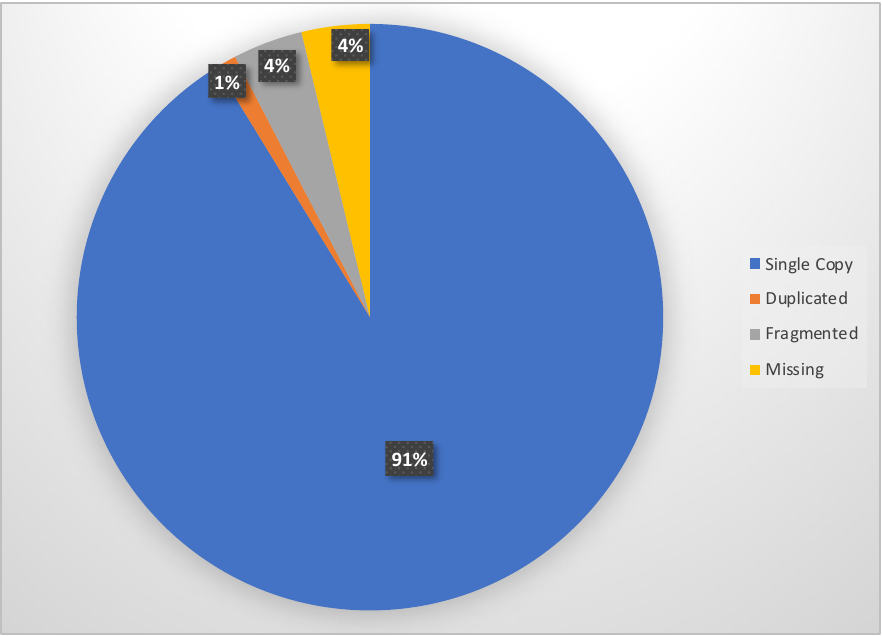

### Chromosome-level assembly

#### Ragout Scaffolding

Lacking long-range sequencing data, we performed the scaffolding of the genome using a reference-assisted approach. Given the availability of multiple reference genomes available, we decided to adopt a multi-reference approach, which might help minimizing the bias introduced by the process. In particular, we decided to use the combination of SibeliaZ v1.1.0 (<https://github.com/medvedevgroup/SibeliaZ/>) for the alignments and Ragout2 (<https://github.com/fenderglass/Ragout/>) for the scaffolding. We used three different references (ARS-UCD1.2, GCA_003369685.2, GCA_003369695.2, which are respectively Hereford, Angus and Brahman). Sequence names have been changed in the UCSC format (>spp.sequenceID; e.g. >hereford.1), matching the input fasta name.

No phylogenetic tree was provided to the software, leaving to Ragout to estimate the relationships from the alignments. The scaffolding has been performed separately for the autosomes, MT, X, Y and the other contigs and scaffolds. The process is represented in the figure below, with the different steps and the genomes used.

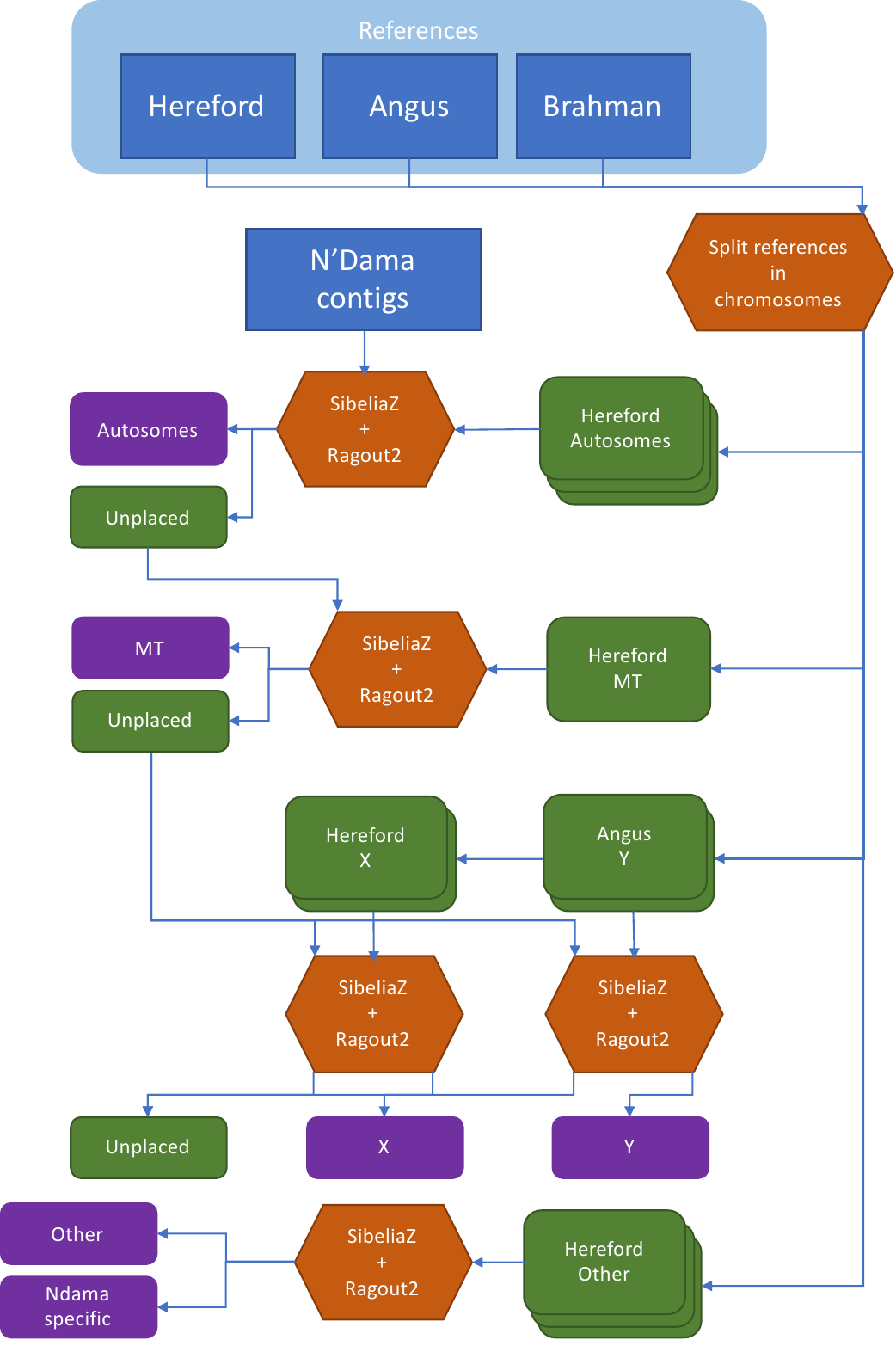

Scheme of the scaffolding step: first, autosomes are scaffolded; the unplaced contigs are first used to create the MT genome, then the remaining used for both X and Y (the same set of contigs have been used to account for pseutoautosomal regions)

SibeliaZ was run with a low k-mer size (-k 21) to use shorter Kmers, slowing down the alignments but increasing the sensitivity.

Most of the contigs are placed on the autosomes, leaving a total sequence length which is roughly the same size as the X chromosome. Worth noticing that one scaffold was identified as on chromosome 7, but the correct position could not be resolved at the time.

| Block | Fragment used | Scfld length | Introduced N | Introduced N (%) | Unplaced Fragments | Unplaced Length | Unplaced Length (%) |
| --- | --- | --- | --- | --- | --- | --- | --- |
| Autosome | 2,081 | 2,504,980,383 | 16,307,546 | 0.65% | 2,564 | 156,203,637 | 5.91% |
| X | 2,194 | 155,751,820 | 64,638,795 | 41.50% | 1,342 | 65,056,028 | 41.66% |
| Y | 640 | 47,396,136 | 8,747,893 | 18.46 | 2,157 | 117,520,810 | 75.25% |
| Mt | 1 | 34,584 | 0 | 0.00 | 2,563 | 156,169,053 | 99.98% |
| Other | 0 | 0 | 0 | 0.00 | 1,177 | 62,647,416 | 100.00% |

The sexual chromosomes present a particularly high number of missing bases, due to the lower coverage derived from the sequencing. Also, the mitogenome, presents twice the expected size. These were considered separately and fixed manually by orientating the fragments and including a gap of ~400 bp.

### Mitochondria misassembly resolution

#### Approach description

Using the QUAST report on the error corrected contigs, it is possible to identify the region of misassembly into the mitochondrial genome.

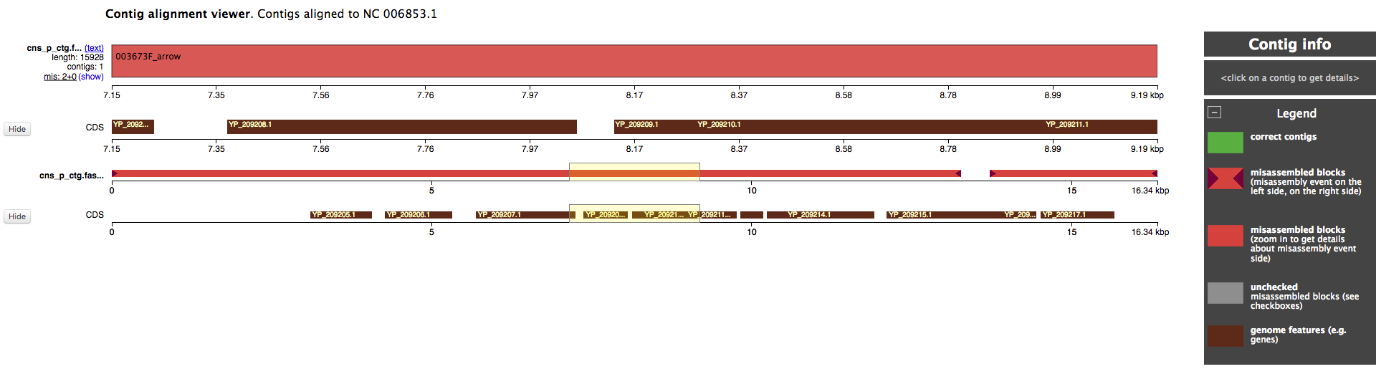

To solve the misassembly, assembled mitochondrial chromosome was aligned to the reference mitochondrial genome using minimap2. Paf alignments were converted to MAF (multiple alignment format). Sorted alignments were then joined manually including a 454 bp gap, reaching the final chromosomal genome length.

#### Scaffold N/L statistics

We calculated some basic statistics of the new chromosome level assembly (Table below). As can be seen, the N50/L50 are high, as expected from a chromosome-level genome, and 90% of the total genome is included in the chromosomes.

| x | Nx | Lx | NGx | LGx | GC% |
| --- | --- | --- | --- | --- | --- |
| 5 | 142,323,786 | 0 | 142,323,786 | 0 | 40.30 |
| 10 | 126,884,692 | 1 | 121,213,004 | 2 | 40.56 |
| 25 | 103,955,525 | 5 | 103,955,525 | 5 | 41.08 |
| 50 | 87,672,696 | 12 | 87,672,696 | 12 | 40.97 |
| 75 | 61,733,321 | 21 | 61,151,627 | 22 | 41.60 |
| 90 | 36,413,080 | 29 | 9,265,313 | 32 | 41.85 |
| 95 | 4,616,499 | 45 | 2,179,503 | 62 | 41.90 |
| 100 | 7,612 | 970 | 7,612 | 970 | 42.00 |

### Gap Filling

We improved the contiguity of the assembly through gap filling using the software LR_GapCloser v1.1. This software split the long reads into 300 bp chunks and map them to the scaffolds using bwa. After that step, it identifies the reads than cover mostly the gap, and use them to fill it. The gap fill stage has been repeated three times. As can be seen in table below, ~50% of the gaps and 34% of Ns have been properly filled:

| Stage | N Gaps | Gaps BP | Bases | Added bp | N50 | L50 |
| --- | --- | --- | --- | --- | --- | --- |
| Initial | 4,885 | 89,694,751 | 2,769,745,704 | 0 | 2,868,616 | 260 |
| Iteration1 | 2,654 | 63,679,091 | 2,769,864,878 | 119,174 | 10,018,179 | 77 |
| Iteration2 | 2,508 | 60,276,639 | 2,769,873,814 | 8,936 | 10,310,270 | 75 |
| Iteration3 | 2,480 | 59,007,146 | 2,769,874,172 | 358 | 10414461 | 74 |

### Genome polishing

Following the gap filling, the genome needs to be finalized through a 5-fold polishing iteration using the short reads and Pilon (v1.23). Illumina short reads (coverage 78X) have been aligned using bwa mem algorithm in 18 chunks, combined with bamtools 2.4.2 and sorted with samtools 1.9.

Below, a table summarising the changes introduced by multiple Pilon runs. Each iteration involved the remapping of the reads to the latest polished version of the assembly.

| Pilon Run | Tot Changes | InDels (<=5bp) | InDels (>5bp) | #SNP | #Gaps fixed |
| --- | --- | --- | --- | --- | --- |
| 1 | 1,967,205 | 1,305,821 | 75,113 | 586,062 | 209 |
| 2 | 268,386 | 107,525 | 37,079 | 123,774 | 8 |
| 3 | 97,875 | 32,851 | 23,049 | 41,973 | 2 |
| 4 | 54,323 | 12,559 | 19,041 | 22,719 | 4 |
| 5 | 38,538 | 6,470 | 17,214 | 14,852 | 2 |

Every iteration reduces the number of changes needed by the assembly. The second Pilon run changed 25% SNPs changed in run 1 (120K vs 580K). The third run changed 33% of SNPs changed from run2 (42K vs 124K), and run 4 changed ~50% of SNPs changed in run3 (22K vs 42K). Similar pattern, but stronger, can be observed with the large indels, that at every iteration decrease massively (from >1,300K to ~12.5K from run 1 to 4). Short indels are less present, and therefore their reduction is less outstanding. However, since small insertions/deletions are a known issue in PacBio sequencing, having a low number of these events confirm that the polishing is proceeding in the right direction. Finally, the gaps fixed changes at every iteration. With the exception of the first, that fixed >200 small gaps, the number of gaps fixed in subsequent iterations depends on how much sequence is added in the previous step. Therefore, it can happen that iteration 4 fix more gaps than iteration 3 (4 vs 2).

### Genome evaluation

Contigs and scaffold metrics for the final assembled genome have been calculated using an in-house script. The quality value (QV) have been calculated using merqury (<https://github.com/marbl/merqury>) on the k-mer counts generated with meryl 1.2 (<https://github.com/marbl/meryl>). QUAST-LG v5 (<http://quast.sourceforge.net>) and FRC_Align (<https://github.com/vezzi/FRC_align>) have been run to assess the genome using a separate reference and an independent evaluation through short reads sequencing. Coverage of the non-N sequence have been calculated with samtools.

|  | N’Dama |
| --- | --- |
| Scaffolds N50 | 104,847,410 |
| Scaffolds L50 | 11 |
| Contigs N50 | 10,726,776 |
| Contigs L50 | 72 |
| Gaps | 2,425 |
| quast genome fract. | 93.9 |
| quast misassemblies | 7,050 |
| Autosomal gaps | 792 |
| QV | 34.3 |
| QV (autosomes) | 37.9 |

The final busco assembly shows 94% of completeness, of which 92.6% in single copy.

| BUSCO | N | %ge |
| --- | --- | --- |
| Complete | 3860 | 94.1% |
| Complete (S) | 3801 | 92.6% |
| Complete (D) | 59 | 1.4% |
| Fragmented | 124 | 3.0% |
| Missing | 120 | 2.9% |
| Total | 4104 | 100.0% |

Coverage plots for the assembly showing the highest level of coverage around the value of 84X (expected = 80X).

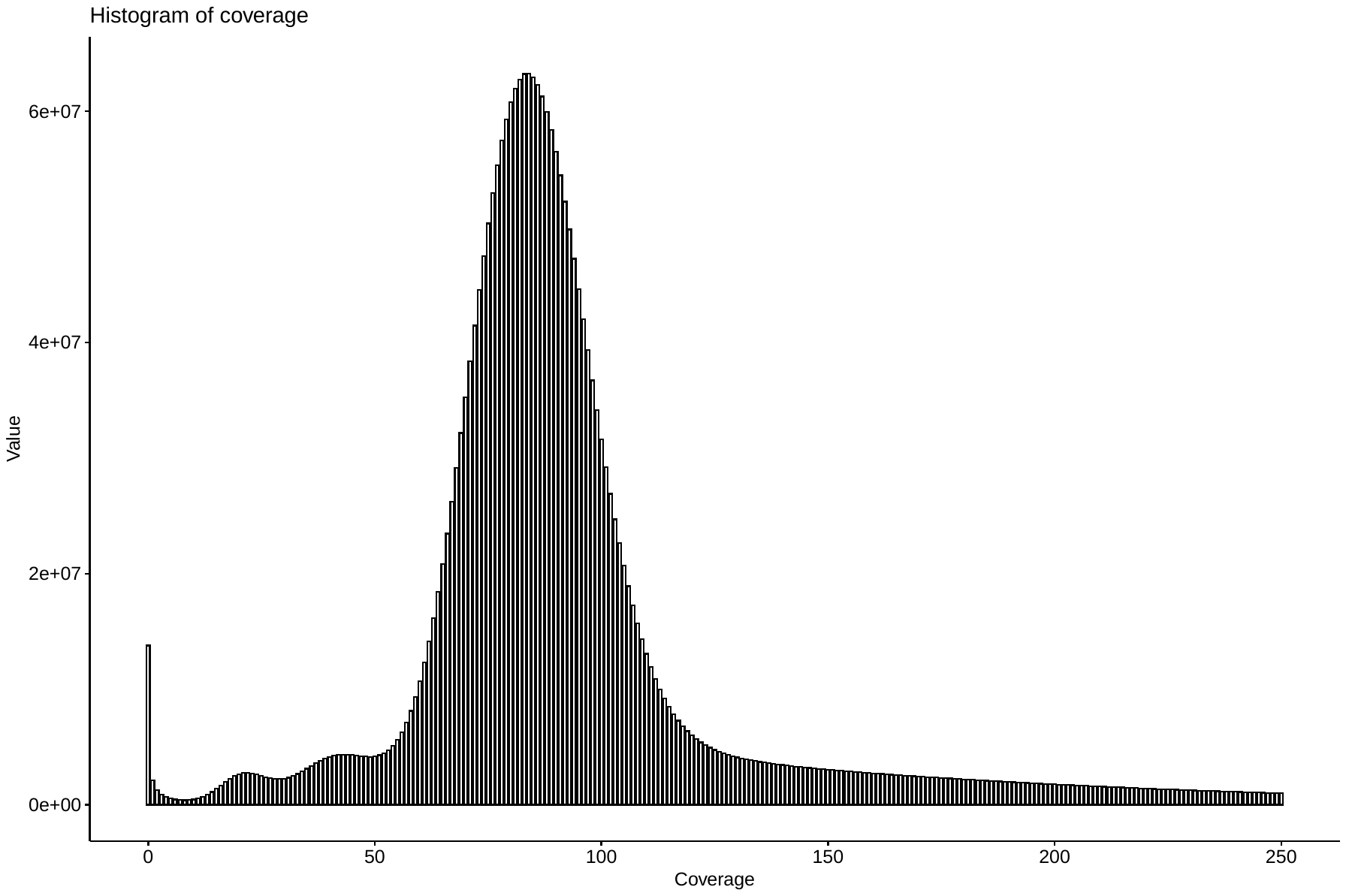

Feature response curve generate through FRC_Align for the assembly is reported below.

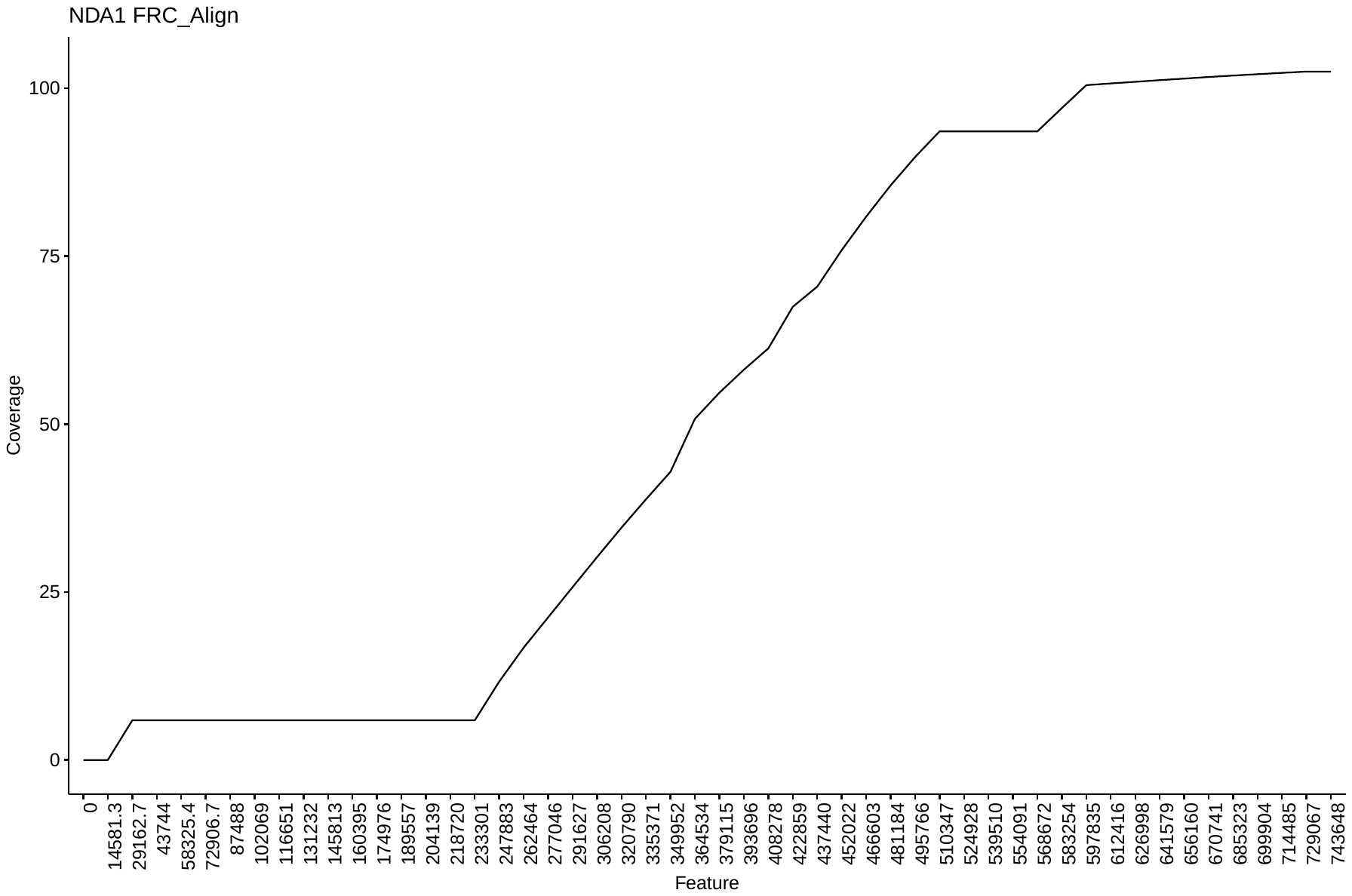

### Repetitive region detection and masking

Following the generation of the genome we performed the masking of the repetitive elements. To identify and mask the repetitive regions, we use a combination of:

1. Dustmasker from NCBI blast+ tool, to mask low-complexity repetitions
2. Windowmasker, to mask interspersed repeats
3. RepeatMasker, that mask interspersed repeats, but that also include trf to mask low-complexity regions

This software generate a bed file with the position of all the repetitive elements in the genome that are then masked using bedtools maskfasta function.

The run of the tools is scattered over multiple jobs, each processing several contigs, to speed up the process. Results from RepeatMasker are then summarised using in-house python script.

### Code availability

All scripts used to generate the assembly are available on GitHub

<https://github.com/evotools/CattleGraphGenomePaper/tree/master/Assembly/NDA1>.
