## Supplementary Note 2 for "A cattle graph genome incorporating global breed diversity"

Supplementary Note 1 – The Ankole assembly report

### Ankole long read sequencing

#### Raw Sequel reads metrics

In table below, there are the metrics for the raw PacBio Sequel subreads.

| Parameter | Value |
| --- | --- |
| #Reads | 9,781,220 |
| Sum | 103,164,533,592 |
| Coverage | 38.21 |
| Minimum | 50 |
| Maximum | 134,087 |
| Mean | 10,547.21 |
| Standard dev. | 8,851.69 |
| Median | 8,448 |
| IQR | 12,813 |

Table 2 – Raw reads base statistics

These parameters confirm the general coverage (~40X) and that the reads seems to have a good median length.

Below, Figure 3 shows the read lengths distribution:

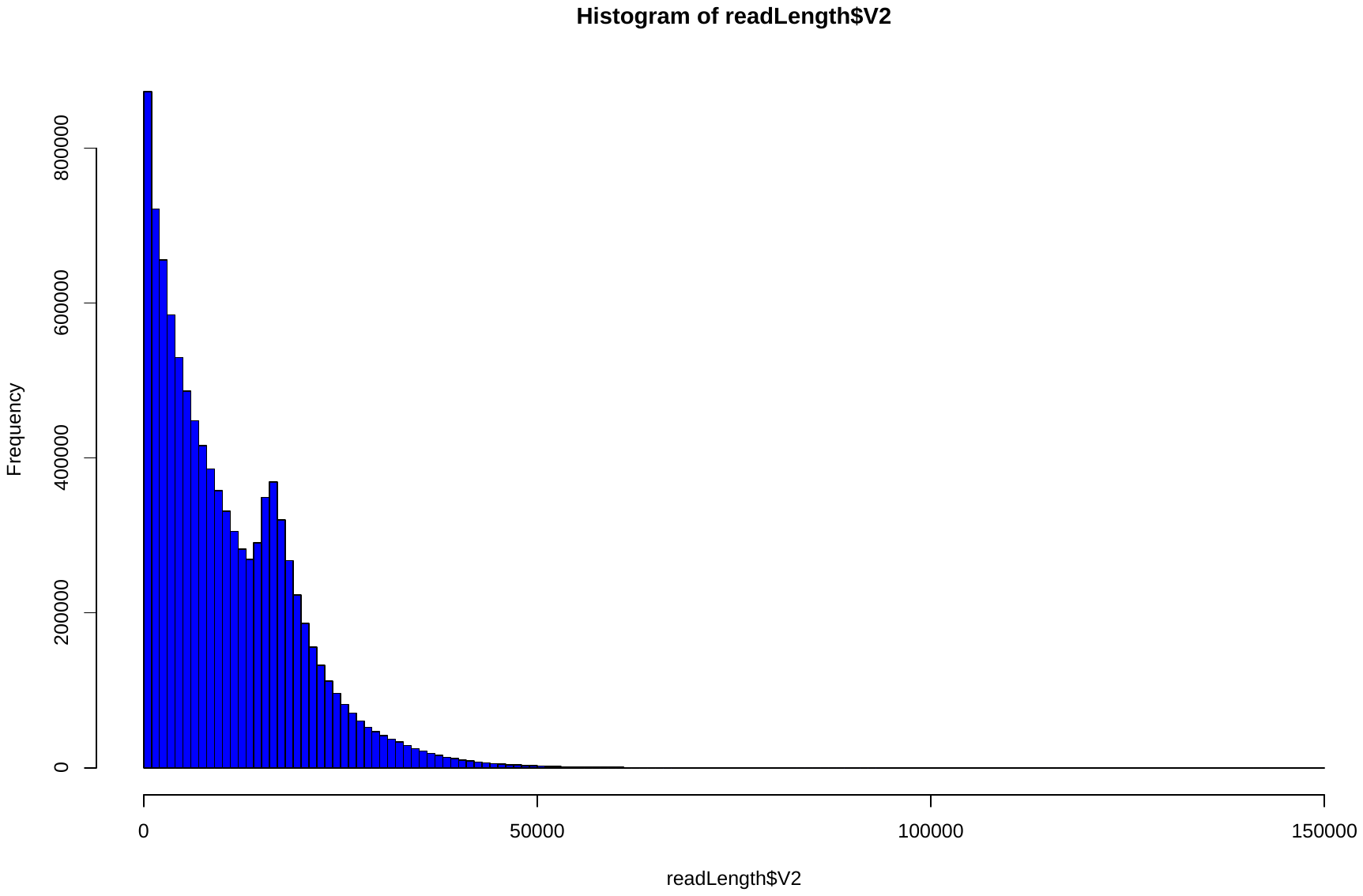

Figure 3 – Read lengths distribution

Analogously to the N’Dama assembly, the reads shows a bimodal distribution, with a first peak to low lengths (<1Kb) and a second at ~17Kb.

### Assembly

We generated two different assemblies using two different assemblers: CANU and WTDBG2. The scripts used to generate the two genomes are reported in GitHub repository. Both assemblies have been polished through WTPOA-cns tool included with the WTDBG2 software. The table below shows the differences between the two assembled genomes.

| Parameter | WTDBG2 | CANU |
| --- | --- | --- |
| N50 | 2,325,045 | 1,800,361 |
| L50 | 329 | 441 |
| BP yielded | 2,703,598,714 | 2,807,643,510 |
| # Sequences | 10,809 | 10,545 |
| BUSCO Complete* | 89.6% | 84.5% |
| BUSCO Fragmented* |  |  |
| BUSCO Missing* |  |  |

Assembler comparison

Both CANU and WTDBG2 provided similar results, with contigs N50 above 1Mb with WTDBG2 showing a slightly higher N50 and lowest L50. Despite that, CANU generated less contigs (300 contigs less) and a longer assembly size (2.8Gb vs 2.7Gb). Most of the statistics still need to be computed for CANU and FALCON, since the two assembler are respectively running the polishing step and generating the contigs.

### Assembly reconciliation tools

Following assembling the genomes, we tried to improve the contiguity of the genome by generating applying a genome reconciliation tool. We used quickmerge, a software that relies on MUMmer alignments filtered for repetitive regions, to detect the overlaps among contigs and combine them into new, longer sequences.

The genomes have been aligned using the nucmer tool in MUMmer4 suite, filtering all the repetitive regions.

The resulting alignments are then used as input for quickmerge, that has been run allowing length cutoff longer than the highest N50 (2.5Mb), considering a minimum alignment length of 50Kb (10X higher than the recommended threshold) and with very stringent cutoffs for the alignments considered for merging (hco > 15, 3X higher than the recommended; c > 5, 3.3X higher than the recommended; values are analogous to (Chakraborty, Baldwin-Brown, Long, & Emerson, 2016)).

|  | Value |
| --- | --- |
| BP yielded | 2,808,308,196 |
| N50 | 8,161,114 |
| L50 | 94 |
| N sequences | 9,270 |
| GC% | 41.99 |

The procedure might have introduced misassemblies, or consolidated some that were present in both assemblies. The use of the Bionano optical mapping will likely fix the largest of these, breaking up chimeric contigs into smaller fragments.

### BioNano optical mapping scaffolding

#### Software description

The generation of scaffolds have been performed using the optical mapping (OM) generated with the BioNano Saphyr machine. Molecules generated have been processed through the BioNano Solve 3.3 pipeline (version 7981).

The optical maps have first been assembled using the quickmerged assembly as reference, and then scaffolded using the hybrid scaffolding pipeline. The workflow introduced 956 cuts to 65 NGS sequences out of 9,271 (0.7% of the total; see table below for output from the conflict solution).

|  | Value |
| --- | --- |
| Number of conflict cuts made to Bionano maps | 170 |
| Number of conflict cuts made to NGS sequences | 956 |
| Number of Bionano maps to be cut | 165 |
| Number of NGS sequences to be cut | 65 |

Eighteen out of the 65 contigs that need to be cut were above 2.5Mb, thirteen between 1Mb and 2.5Mb and 31 below 1Mb, with all contigs that had more than 5 cuts below the 1Mb threshold, suggesting the presence of misassemblies at the assembler level. More than 60% of the contigs (40/65) were split in two fragments, 24% in three fragments and the 15% in >4 fragments.

| Ctg ID | CTG Size | # Fragments | Smaller Fragment Size | Larger Fragment Size | All Fragment Sizes |
| --- | --- | --- | --- | --- | --- |
| 6399 | 235,114 | 10 | 2608 | 120196 | 41052;7461;13523;12100;4012;2608;5415;26130;2608;120196 |
| 3144 | 163,846 | 9 | 2559 | 115541 | 28708;2559;4002;2604;2608;2605;2605;2606;115541 |
| 2772 | 371,428 | 7 | 59 | 158049 | 50584;40337;59;80208;17888;24297;158049 |
| 3438 | 113,603 | 7 | 3989 | 29329 | 27292;3989;7749;18511;22719;4008;29329 |
| 637 | 794,184 | 5 | 59 | 398055 | 163493;4325;228248;59;398055 |
| 1731 | 887,286 | 5 | 21386 | 443612 | 82670;79794;443612;21386;259820 |
| 1915 | 811,417 | 4 | 10574 | 559832 | 559832;10574;10846;230162 |
| 9129 | 12,039,443 | 4 | 59 | 8774036 | 3237364;27981;59;8774036 |
| 9247 | 12,474,065 | 4 | 8208 | 12092034 | 12092034;48131;8208;325689 |
| 1147 | 1,585,356 | 3 | 23507 | 1206018 | 355829;23507;1206018 |
| 1270 | 1,331,647 | 3 | 6379 | 732968 | 732968;6379;592298 |
| 2109 | 322,094 | 3 | 22980 | 245702 | 53410;22980;245702 |
| 2157 | 618,540 | 3 | 13853 | 568760 | 35925;13853;568760 |
| 2322 | 453,240 | 3 | 9931 | 302333 | 302333;9931;140974 |
| 3383 | 109,369 | 3 | 2608 | 76059 | 30700;2608;76059 |
| 3428 | 142,634 | 3 | 4012 | 94822 | 43798;4012;94822 |
| 5960 | 2,404,879 | 3 | 6436 | 2001399 | 2001399;6436;397042 |
| 6007 | 609,323 | 3 | 39787 | 490829 | 78705;39787;490829 |
| 6237 | 256,704 | 3 | 10727 | 161749 | 161749;10727;84226 |
| 6413 | 88,473 | 3 | 4419 | 58615 | 58615;4419;25437 |
| 8727 | 1,889,629 | 3 | 23394 | 1472041 | 394192;23394;1472041 |
| 9094 | 50,349,372 | 3 | 6311 | 28628641 | 21714418;6311;28628641 |
| 9143 | 21,002,380 | 3 | 32861 | 13946044 | 13946044;32861;7023473 |
| 9194 | 2,812,833 | 3 | 28719 | 2686466 | 97646;28719;2686466 |
| 9229 | 8,027,832 | 3 | 59 | 4443568 | 3584203;59;4443568 |
| 8 | 4,496,967 | 2 | 1853501 | 2643465 | 1853501;2643465 |
| 72 | 181,619 | 2 | 32189 | 149429 | 32189;149429 |
| 216 | 182,234 | 2 | 46042 | 136191 | 46042;136191 |
| 594 | 2,496,872 | 2 | 29398 | 2467473 | 29398;2467473 |
| 629 | 1,263,114 | 2 | 23285 | 1239828 | 1239828;23285 |
| 800 | 1,114,663 | 2 | 445969 | 668693 | 445969;668693 |
| 972 | 1,878,577 | 2 | 235915 | 1642661 | 235915;1642661 |
| 1084 | 1,568,556 | 2 | 730844 | 837711 | 730844;837711 |
| 1312 | 1,409,293 | 2 | 104420 | 1304872 | 104420;1304872 |
| 1412 | 1,204,556 | 2 | 181058 | 1023497 | 1023497;181058 |
| 1868 | 876,811 | 2 | 106805 | 770005 | 770005;106805 |
| 2130 | 445,190 | 2 | 119237 | 325952 | 119237;325952 |
| 2447 | 381,485 | 2 | 90006 | 291478 | 291478;90006 |
| 2809 | 122,592 | 2 | 36774 | 85817 | 85817;36774 |
| 2968 | 122,937 | 2 | 43020 | 79916 | 79916;43020 |
| 3311 | 141,503 | 2 | 32150 | 109352 | 32150;109352 |
| 5993 | 1,182,297 | 2 | 64329 | 1117967 | 64329;1117967 |
| 6037 | 2,348,223 | 2 | 133913 | 2214309 | 2214309;133913 |
| 6058 | 245,737 | 2 | 60375 | 185361 | 60375;185361 |
| 6143 | 288,439 | 2 | 135856 | 152582 | 152582;135856 |
| 6168 | 508,293 | 2 | 51190 | 457102 | 457102;51190 |
| 6230 | 270,168 | 2 | 75577 | 194590 | 194590;75577 |
| 6251 | 210,208 | 2 | 98570 | 111637 | 98570;111637 |
| 6253 | 332,684 | 2 | 37785 | 294898 | 37785;294898 |
| 6256 | 405,643 | 2 | 36263 | 369379 | 36263;369379 |
| 6360 | 357,635 | 2 | 117202 | 240432 | 240432;117202 |
| 9103 | 13,274,084 | 2 | 54086 | 13219997 | 13219997;54086 |
| 9126 | 5,989,548 | 2 | 391061 | 5598486 | 391061;5598486 |
| 9133 | 5,315,080 | 2 | 136169 | 5178910 | 136169;5178910 |
| 9167 | 13,510,537 | 2 | 158305 | 13352231 | 158305;13352231 |
| 9182 | 5,134,164 | 2 | 1835694 | 3298469 | 3298469;1835694 |
| 9189 | 10,293,330 | 2 | 1722992 | 8570337 | 8570337;1722992 |
| 9193 | 6,841,681 | 2 | 102276 | 6739404 | 102276;6739404 |
| 9204 | 8,229,963 | 2 | 2445330 | 5784632 | 2445330;5784632 |
| 9206 | 4,142,798 | 2 | 1118188 | 3024609 | 3024609;1118188 |
| 9211 | 6,756,289 | 2 | 1091867 | 5664421 | 5664421;1091867 |
| 9219 | 9,152,709 | 2 | 742149 | 8410559 | 8410559;742149 |
| 9241 | 16,694,204 | 2 | 1420161 | 15274042 | 1420161;15274042 |
| 9252 | 8,673,619 | 2 | 254878 | 8418740 | 254878;8418740 |
| 9265 | 11,484,869 | 2 | 74937 | 11409931 | 74937;11409931 |

The resulting scaffolds, in comparison with the contig level assemblies, show the following metrics:

| Assembly | Bp | #Scf | scfN50 (Mb) | #Ctg | ctgN50 | ctgL50 | #Ns | #Gaps | LongestGap |
| --- | --- | --- | --- | --- | --- | --- | --- | --- | --- |
| WTDBG2 | 2,703,598,714 | NA | NA | 10,809 | 2,325,045 | 329 | NA | NA | NA |
| CANU | 2,807,643,510 | NA | NA | 10,545 | 1,800,361 | 441 | NA | NA | NA |
| QUICKMERGE | 2,808,308,196 | NA | NA | 9,270 | 8,161,114 | 94 | NA | NA | NA |
| QKM_BNSCF | 2,808,308,196 | 7,581 | 85.414 | 9,388 | 7,823,042 | 97 | 118,404,067 | 1,807 | 5,536,000 |

The next stage will be the gap filling and polishing of the genome generated from the Solve pipeline.

### Gap Filling

Following the definition of the scaffolded version of the genome, the next step to be performed is the gap filling in order to increase the contiguity of the assembly. We used the scaffolds generated starting from the quickmerge contigs and the raw PacBio long reads. The software used to perform this step is LR_GapCloser (Xu et al., 2018). The following table shows the results of each of the 3-fold gap-filling iterations, with the sequential improvements:

| Assembly | Bp | #Scf | scfN50 | #Ctg | ctgN50 | ctgL50 | #Ns | #Gaps | LongestGap |
| --- | --- | --- | --- | --- | --- | --- | --- | --- | --- |
| WTDBG2 | 2,703,598,714 | NA | NA | 10,809 | 2,325,045 | 329 | NA | NA | NA |
| CANU | 2,807,643,510 | NA | NA | 10,545 | 1,800,361 | 441 | NA | NA | NA |
| QUICKMERGE | 2,808,308,196 | NA | NA | 9,270 | 8,161,114 | 94 | NA | NA | NA |
| QKM_BNSCF | 2,808,417,765 | 7,581 |  | 9,388 | 7,823,042 | 97 | 118,407,067 | 1,807 | 5,536,000 |
| QKM_BNSCF  LRGC (I1) | 2,816,250,667 | 7,581 |  | 8,822 | 7,823,042 | 52 | 98,366,957 | 1,241 | 5,508,511 |
| QKM_BNSCF  LRGC (I2) | 2,817,716,955 | 7,581 |  | 8,575 | 16,946,366 | 49 | 90,935,046 | 994 | 5,508,511 |
| QKM_BNSCF  LRGC (I3) | 2,818,214,162 | 7,581 |  | 8,505 | 17,812,669 | 49 | 86,562,567 | 924 | 5,508,511 |

Next step in this analysis involves the polishing of the scaffolds, using Illumina short-reads, to further improve the overall quality of the genome.

### Genome polishing

Following the gap filling, the genome was finalized through 5-fold iterations of polishing through Pilon v1.23 with 78X Illumina short reads mapped scaffolded, gap-filled genome. Illumina short reads have then been aligned through bwa mem algorithm in 18 chunks, joined with bamtools 2.4.2 and sorted with samtools 1.9.

Below, a table summarising the changes introduced by multiple Pilon runs. Each iteration involved the remapping of the reads to the latest polished version of the assembly.

| Pilon Run | Tot Changes | InDels (<=5bp) | InDels (>5bp) | #SNP | #Gaps fixed |
| --- | --- | --- | --- | --- | --- |
| 1 | 4,099,358 | 3,385,056 | 135,069 | 579,233 | 7 |
| 2 | 428,490 | 207,288 | 66,250 | 154,952 | 8 |
| 3 | 155,155 | 72,104 | 31,409 | 51,642 | 4 |
| 4 | 75,285 | 23,964 | 26,429 | 24,892 | 1 |
| 5 | 50,588 | 12,951 | 23,749 | 13,888 | 1 |

Every iteration reduces the number of changes needed by the assembly. The number of gaps fixed in each iteration varies depending on the added sequence from the previous iteration.

### Chromosome assignment

#### Alignments and assignments

Following completion of the assembly, we tried to identify which scaffolds corresponded to the autosomes, sexual chromosomes and mitogenome. To do so, we first aligned the scaffolds to the 1000 bull reference genome using minimap2. The resulting paf were then processed through a custom R scripts to extract the alignments that better suited each autosome. The result is reported in the table below, with most of the chromosomes corresponding to a single scaffold, with high percentage of identity:

| **Query** | **Target** | **Aligned** | **Tgt Length** | **Qry Length** | **Ratio Qry aligned** | **Ratio Tgt aligned** |
| --- | --- | --- | --- | --- | --- | --- |
| Super-Scaffold_100001 | 1 | 155901980 | 158534110 | 156527526 | 0.9833971 | 0.9960036 |
| Super-Scaffold_100002 | 2 | 137697348 | 136231102 | 138302011 | 1.0107629 | 0.995628 |
| Super-Scaffold_100003 | 3 | 119614633 | 121005158 | 120813661 | 0.9885085 | 0.9900754 |
| Super-Scaffold_100005 | 4 | 118799771 | 120000601 | 119914157 | 0.9899931 | 0.9907068 |
| Super-Scaffold_100004 | 5 | 116688774 | 120089316 | 122124786 | 0.9716832 | 0.9554881 |
| Super-Scaffold_100007 | 6 | 111516922 | 117806340 | 112516630 | 0.9466122 | 0.991115 |
| Super-Scaffold_100008 | 7 | 109482276 | 110682743 | 110533585 | 0.989154 | 0.9904888 |
| Super-Scaffold_100006 | 8 | 112418051 | 113319770 | 114147848 | 0.9920427 | 0.984846 |
| Super-Scaffold_100010 | 9 | 104081408 | 105454467 | 104606559 | 0.9869796 | 0.9949798 |
| Super-Scaffold_100011 | 10 | 100737099 | 103308737 | 103296156 | 0.9751073 | 0.975226 |
| Super-Scaffold_100009 | 11 | 105979159 | 106982474 | 107065538 | 0.9906217 | 0.9898531 |
| Super-Scaffold_100012 | 12 | 83734362 | 87216183 | 89974165 | 0.9600783 | 0.9306489 |
| Super-Scaffold_9109 | 13 | 82263160 | 83472345 | 83163653 | 0.9855139 | 0.989172 |
| Super-Scaffold_9214 | 14 | 79091931 | 82403003 | 82216954 | 0.9598186 | 0.9619905 |
| Super-Scaffold_100013 | 15 | 81771760 | 85007780 | 84415766 | 0.9619327 | 0.9686788 |
| Super-Scaffold_100016 | 16 | 73121424 | 81013979 | 74962467 | 0.9025779 | 0.9754405 |
| Super-Scaffold_100017 | 17 | 72180629 | 73167244 | 72786150 | 0.9865156 | 0.9916808 |
| Super-Scaffold_9192 | 18 | 65305312 | 65820629 | 67291221 | 0.9921709 | 0.9704878 |
| Super-Scaffold_100021 | 19 | 62919788 | 63449741 | 63302509 | 0.9916477 | 0.9939541 |
| Super-Scaffold_100018 | 20 | 71188688 | 71974595 | 72406059 | 0.9890808 | 0.9831869 |
| Super-Scaffold_100019 | 21 | 68731579 | 69862954 | 70766496 | 0.9838058 | 0.9712446 |
| Super-Scaffold_100023 | 22 | 59638881 | 60773035 | 61182121 | 0.9813379 | 0.9747763 |
| Super-Scaffold_100024 | 23 | 50754560 | 52498615 | 52932239 | 0.966779 | 0.9588591 |
| Super-Scaffold_100022 | 24 | 62068383 | 62317253 | 62232639 | 0.9960064 | 0.9973606 |
| Super-Scaffold_100030 | 25 | 42510937 | 42350435 | 42999529 | 1.0037899 | 0.9886373 |
| Super-Scaffold_100025 | 26 | 50788676 | 51992305 | 51362451 | 0.9768499 | 0.9888289 |
| Super-Scaffold_100028 | 27 | 43889740 | 45612108 | 45509546 | 0.9622388 | 0.9644073 |
| Super-Scaffold_100029 | 28 | 37006736 | 45940150 | 42633662 | 0.8055423 | 0.8680168 |
| Super-Scaffold_100026 | 29 | 51374455 | 51098607 | 53282723 | 1.0053983 | 0.964186 |
| tig00008859_obj | MT | 35866 | 16340 | 36534 | 2.1949816 | 0.9817157 |
| tig00009004_obj | MT | 16102 | 16340 | 17364 | 0.9854345 | 0.9273209 |
| tig00009055_obj | MT | 16988 | 16340 | 17519 | 1.0396573 | 0.9696901 |
| tig00009122_obj | MT | 14966 | 16340 | 16578 | 0.9159119 | 0.9027627 |
| tig00009170_obj | MT | 15444 | 16340 | 17012 | 0.9451652 | 0.9078298 |
| tig00009193_obj | MT | 13312 | 16340 | 16003 | 0.8146879 | 0.831844 |
| tig00009207_obj | MT | 13443 | 16340 | 14566 | 0.822705 | 0.9229027 |
| tig00009254_obj | MT | 14195 | 16340 | 15426 | 0.8687271 | 0.9201997 |

The lower identity of chromosome 28 can be linked to the presence of a large gap (>5Mb) reducing the total alignments percentage. Mitogenome has been assembled multiple times in several contigs. The one closer in size to the actual mitogenome is *tig00009055_obj*, which we suggest as the actual mitochondrial genome. Finally, both X and Y have not been properly scaffolded. They are covered by a various number of scaffolds and/or contigs, and never reaching the total expected chromosomal length.

### Genome evaluation

Contigs and scaffold metrics for the final assembled genome have been calculated using an in-house script. The quality value (QV) have been calculated using merqury (<https://github.com/marbl/merqury>) on the k-mer counts generated with meryl 1.2 (<https://github.com/marbl/meryl>). QUAST-LG v5 (<http://quast.sourceforge.net>) and FRC_Align (<https://github.com/vezzi/FRC_align>) have been run to assess the genome using a separate reference and an independent evaluation through short reads sequencing. Coverage of the non-N sequence have been calculated with samtools.

|  | Ankole |
| --- | --- |
| Scaffolds N50 | 84,415,766 |
| Scaffolds L50 | 12 |
| Contigs N50 | 18,610,934 |
| Contigs L50 | 49 |
| Gaps | 904 |
| Autosomal gaps | 296 |
| QUAST genome fract. | 94.0% |
| QUAST misassemblies | 5,907 |
| QV | 30.6 |
| QV (autosomes) | 34.2 |

The final busco assembly shows 93% of completeness, of which 91.0% in single copy.

| BUSCO | N | %ge |
| --- | --- | --- |
| Complete | 3819 | 93.1% |
| Complete (Single) | 3733 | 91.0% |
| Complete (Duplicate) | 86 | 2.1% |
| Fragmented | 125 | 3.0% |
| Missing | 160 | 3.9% |
| Total | 4104 | 100.0% |

The coverage plot shows the highest coverage at 89X.

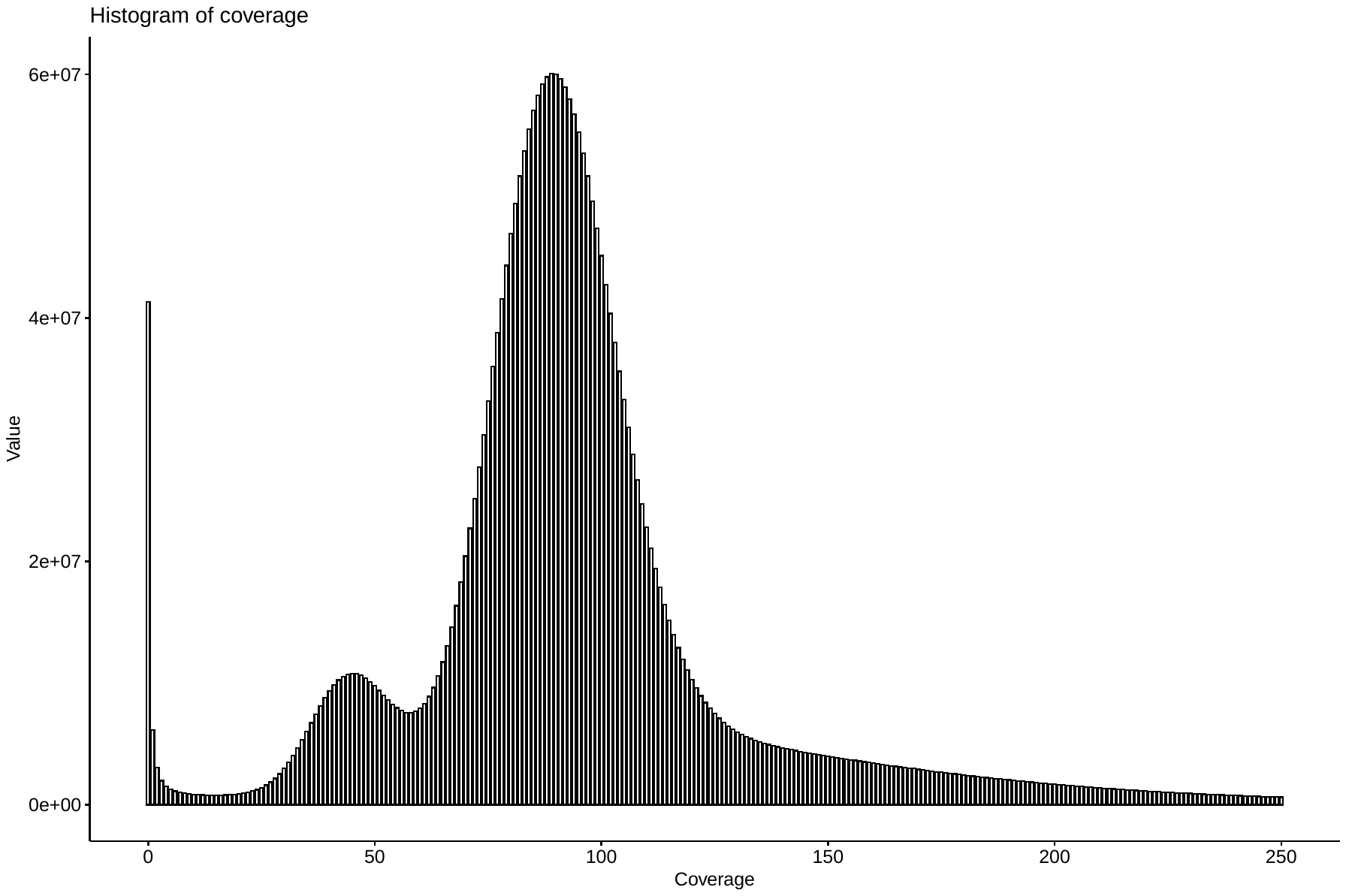

The unusual increase in lower coverage is due to the sparse unplaced contigs, and when removed the second peak almost disappear:

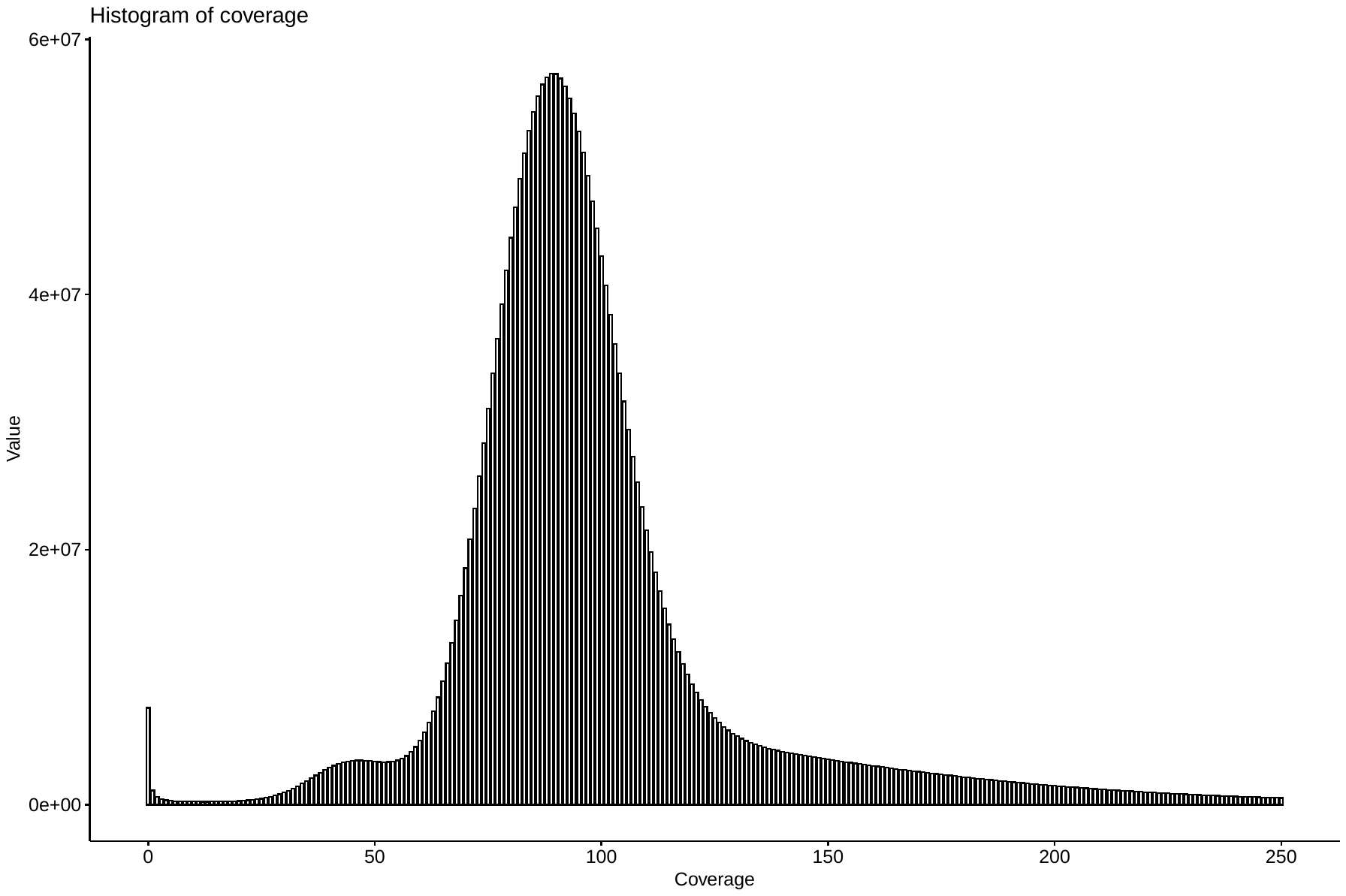

The Feature Response Curve generated with FRC_Align is reported below.

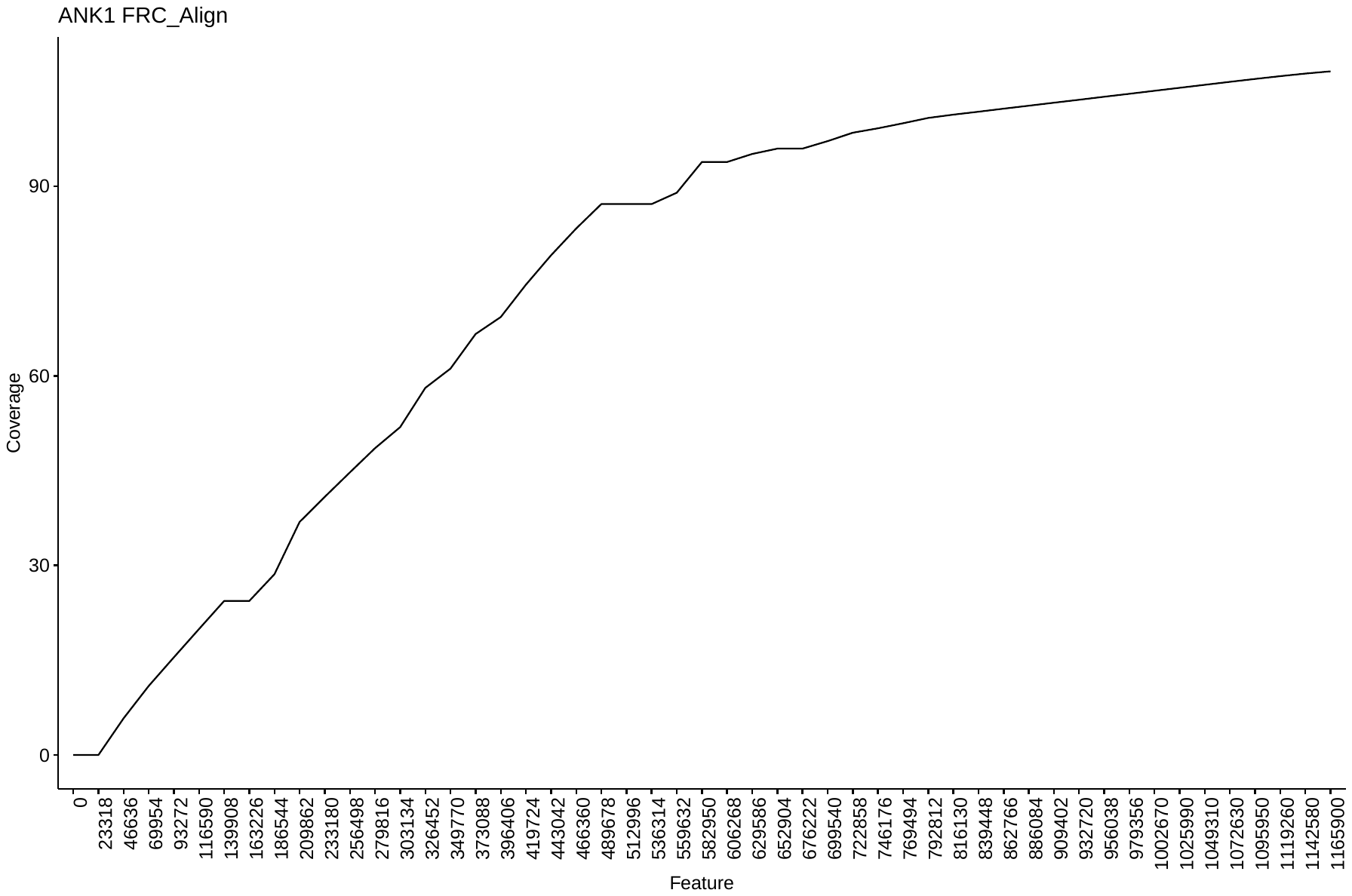

### Repetitive region detection and masking

Following the generation of the genome we performed the masking of the repetitive elements. To identify and mask the repetitive regions, we use a combination of:

1. Dustmasker from NCBI blast+ tool, to mask low-complexity repetitions
2. Windowmasker, to mask interspersed repeats
3. RepeatMasker, that mask interspersed repeats, but that also include trf to mask low-complexity regions

This software generate a bed file with the position of all the repetitive elements in the genome that are then masked using bedtools maskfasta function.

The run of the tools is scattered over multiple jobs, each processing several contigs, to speed up the process. Results from RepeatMasker are then summarised using in-house python script.

### Code availability

All scripts used to generate the assembly are available on GitHub

<https://github.com/evotools/CattleGraphGenomePaper/tree/master/Assembly/ANK1>
