## Supplementary Note 3 for "A cattle graph genome incorporating global breed diversity"

### Supplementary Note 3 – Comparison of variants by algorithm

##### Number of variants

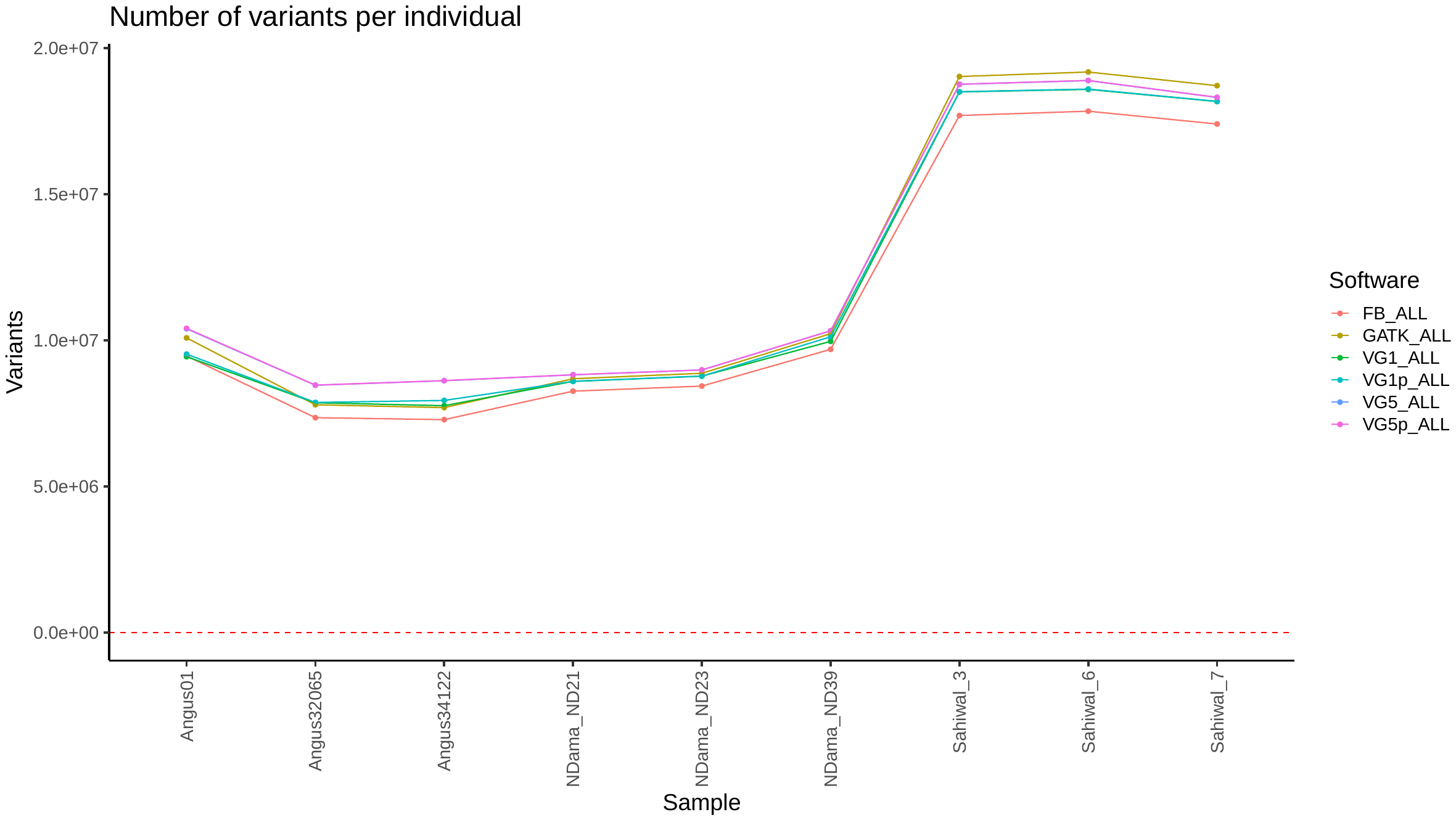

##### Variant number by individual and algorithm

| **ALGORITHM** | **SIZE** | **Angus** | | | **N'Dama** | | | **Sahiwal** | | |
| --- | --- | --- | --- | --- | --- | --- | --- | --- | --- | --- |
|  |  | **Angus01** | **Angus32065** | **Angus34122** | **NDama_ND21** | **NDama_ND23** | **NDama_ND39** | **Sahiwal_3** | **Sahiwal_6** | **Sahiwal_7** |
| FB | 0 | 9,433,407 | 8,680,825 | 8,647,275 | 10,293,619 | 10,427,815 | 11,365,489 | 22,267,905 | 21,961,629 | 21,134,702 |
|  | 30 | 1,247,675 | 1,291,245 | 1,286,982 | 1,459,222 | 1,465,023 | 1,528,898 | 2,601,979 | 2,563,991 | 2,465,586 |
|  | 100 | 699 | 978 | 912 | 5,096 | 5,147 | 5,258 | 11,094 | 10,589 | 10,292 |
|  | 500 | 0 | 0 | 0 | 0 | 0 | 0 | 0 | 0 | 0 |
|  | 1000 | 0 | 0 | 0 | 0 | 0 | 0 | 0 | 0 | 0 |
|  | 5000 | 0 | 0 | 0 | 0 | 0 | 0 | 0 | 0 | 0 |
|  | >=5001 | 0 | 0 | 0 | 0 | 0 | 0 | 0 | 0 | 0 |
| GATK | 0 | 9,714,748 | 9,122,613 | 9,062,461 | 10,781,189 | 10,925,493 | 11,938,304 | 23,933,608 | 23,600,031 | 22,705,447 |
|  | 30 | 1,410,833 | 1,390,929 | 1,365,907 | 1,648,530 | 1,658,077 | 1,741,077 | 2,916,412 | 2,880,400 | 2,791,150 |
|  | 100 | 11,414 | 13,255 | 12,474 | 18,693 | 18,947 | 20,196 | 38,075 | 37,585 | 36,361 |
|  | 500 | 598 | 829 | 731 | 2,040 | 2,007 | 2,082 | 3,840 | 3,609 | 3,593 |
|  | 1000 | 0 | 0 | 0 | 0 | 0 | 0 | 0 | 0 | 0 |
|  | 5000 | 0 | 0 | 0 | 0 | 0 | 0 | 0 | 0 | 0 |
|  | >=5001 | 0 | 0 | 0 | 0 | 0 | 0 | 0 | 0 | 0 |
| VG1 | 0 | 9,310,979 | 9,146,153 | 8,982,935 | 10,637,904 | 10,779,804 | 11,644,564 | 23,176,834 | 22,793,550 | 21,995,140 |
|  | 30 | 1,202,143 | 1,200,398 | 1,180,741 | 1,403,965 | 1,413,178 | 1,499,336 | 2,515,032 | 2,495,894 | 2,409,959 |
|  | 100 | 503 | 753 | 623 | 3,521 | 3,587 | 3,611 | 7,785 | 7,263 | 7,530 |
|  | 500 | 0 | 0 | 0 | 0 | 0 | 0 | 0 | 0 | 0 |
|  | 1000 | 0 | 0 | 0 | 0 | 0 | 0 | 0 | 0 | 0 |
|  | 5000 | 0 | 0 | 0 | 0 | 0 | 0 | 0 | 0 | 0 |
|  | >=5001 | 0 | 0 | 0 | 0 | 0 | 0 | 0 | 0 | 0 |
| VG1P | 0 | 9,426,559 | 9,161,053 | 9,195,152 | 10,642,495 | 10,782,819 | 11,803,691 | 23,192,625 | 22,817,076 | 22,009,622 |
|  | 30 | 1,202,013 | 1,203,250 | 1,207,983 | 1,406,005 | 1,415,432 | 1,504,931 | 2,519,744 | 2,500,724 | 2,413,828 |
|  | 100 | 552 | 814 | 685 | 3,580 | 3,620 | 3,851 | 7,940 | 7,421 | 7,664 |
|  | 500 | 8 | 8 | 3 | 6 | 6 | 5 | 12 | 13 | 12 |
|  | 1000 | 0 | 0 | 0 | 0 | 0 | 0 | 0 | 0 | 0 |
|  | 5000 | 0 | 0 | 0 | 0 | 0 | 0 | 0 | 0 | 0 |
|  | >=5001 | 0 | 0 | 0 | 0 | 0 | 0 | 0 | 0 | 0 |
| VG5 | 0 | 10,384,667 | 9,830,558 | 9,956,794 | 10,963,951 | 11,081,224 | 12,099,440 | 23,604,880 | 23,263,468 | 22,289,674 |
|  | 30 | 1,362,140 | 1,350,017 | 1,363,988 | 1,564,503 | 1,575,635 | 1,683,092 | 2,786,138 | 2,767,011 | 2,650,831 |
|  | 100 | 14,155 | 14,868 | 14,089 | 17,711 | 17,992 | 19,057 | 32,716 | 32,308 | 31,196 |
|  | 500 | 8,947 | 8,943 | 8,665 | 10,028 | 10,119 | 10,739 | 16,620 | 16,619 | 16,134 |
|  | 1000 | 2,632 | 2,510 | 2,475 | 2,607 | 2,610 | 2,858 | 4,279 | 4,208 | 4,089 |
|  | 5000 | 3,357 | 3,266 | 3,227 | 3,317 | 3,268 | 3,550 | 4,981 | 4,943 | 4,871 |
|  | >=5001 | 343 | 340 | 341 | 387 | 359 | 382 | 447 | 436 | 443 |
| VG5P | 0 | 10,404,632 | 9,834,954 | 9,961,919 | 10,968,870 | 11,084,581 | 12,101,691 | 23,613,489 | 23,274,904 | 22,299,958 |
|  | 30 | 1,361,051 | 1,349,330 | 1,362,827 | 1,563,403 | 1,574,625 | 1,681,623 | 2,782,881 | 2,764,108 | 2,647,688 |
|  | 100 | 14,128 | 14,865 | 14,117 | 17,731 | 17,984 | 19,023 | 32,740 | 32,317 | 31,201 |
|  | 500 | 8,962 | 8,941 | 8,655 | 10,030 | 10,105 | 10,724 | 16,558 | 16,580 | 16,095 |
|  | 1000 | 2,619 | 2,506 | 2,475 | 2,617 | 2,605 | 2,853 | 4,259 | 4,187 | 4,085 |
|  | 5000 | 3,368 | 3,230 | 3,218 | 3,299 | 3,268 | 3,542 | 4,931 | 4,939 | 4,834 |
|  | >=5001 | 345 | 347 | 341 | 386 | 356 | 377 | 449 | 437 | 444 |

#### Scenario A: 11M variants as known

##### Variant number by size

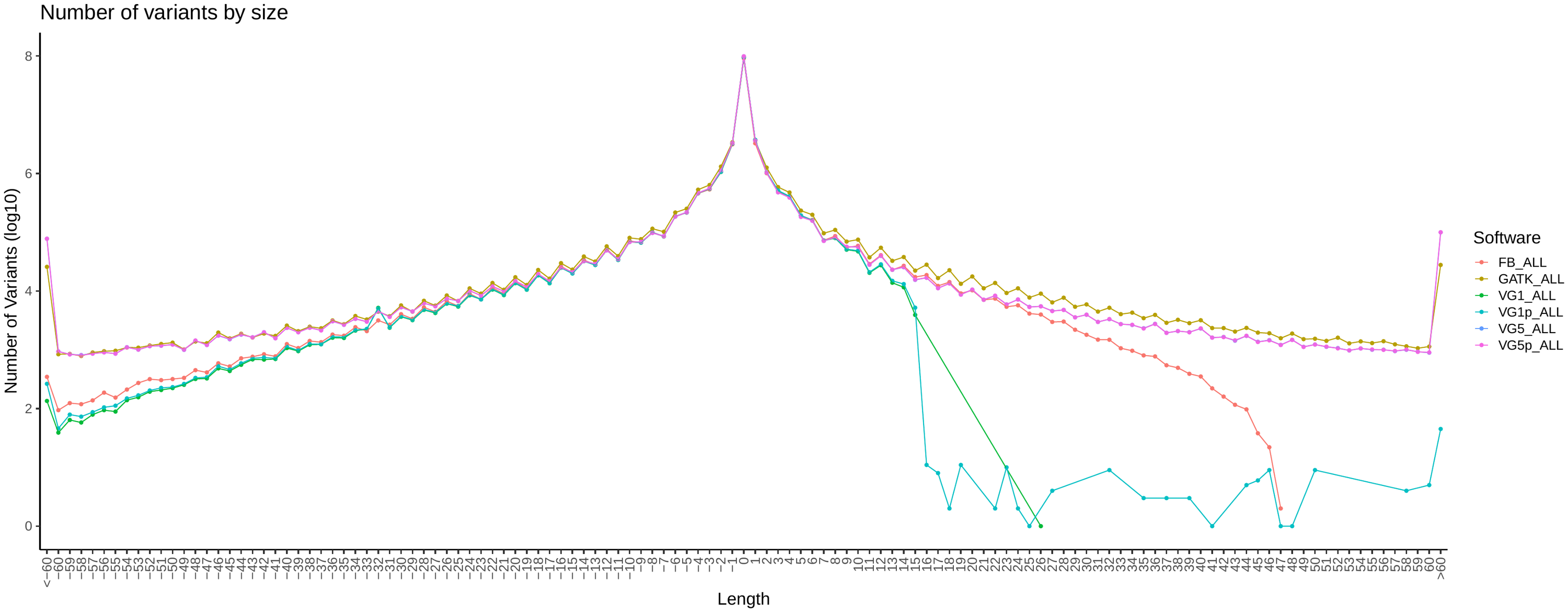

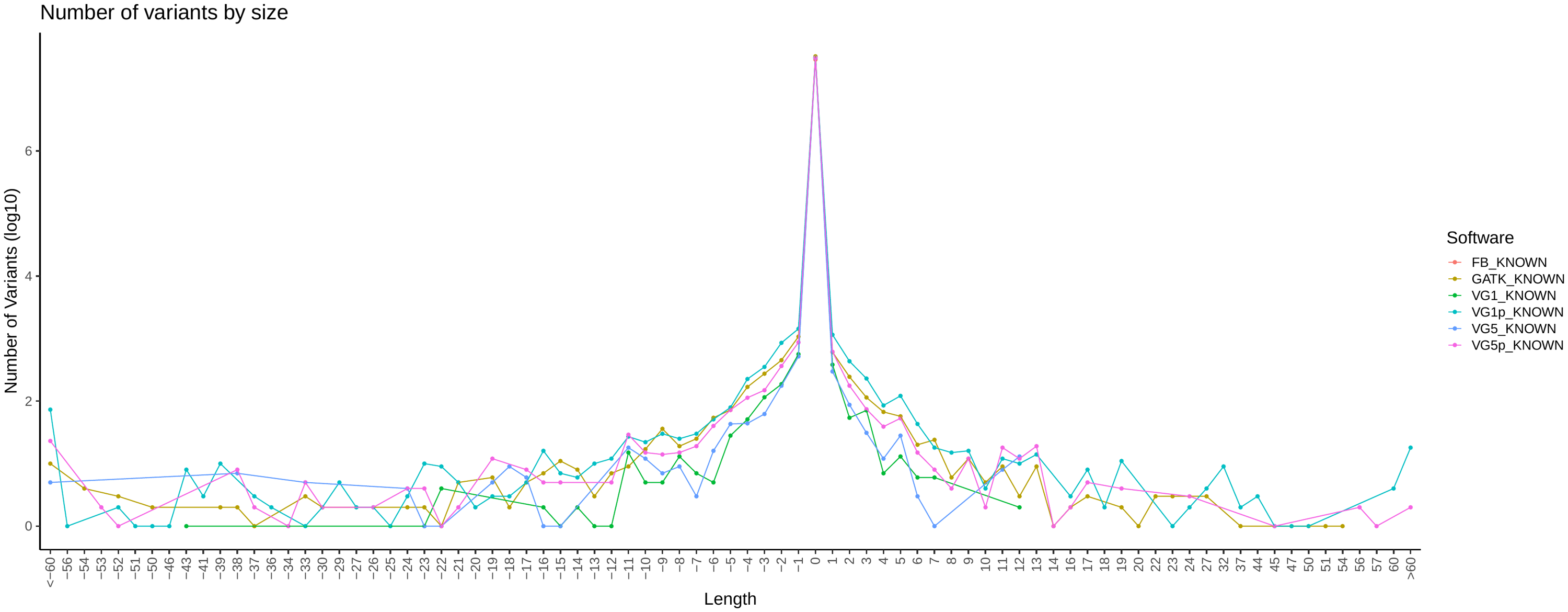

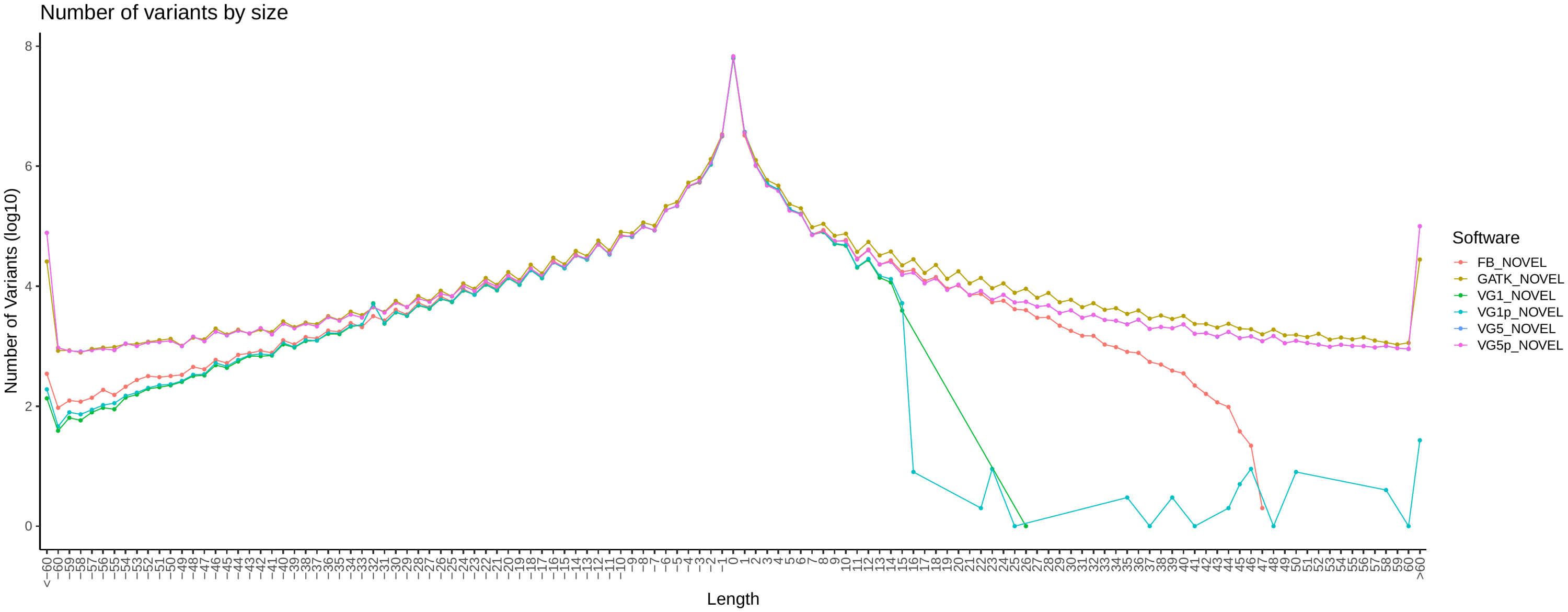

##### Variant QUAL by Size

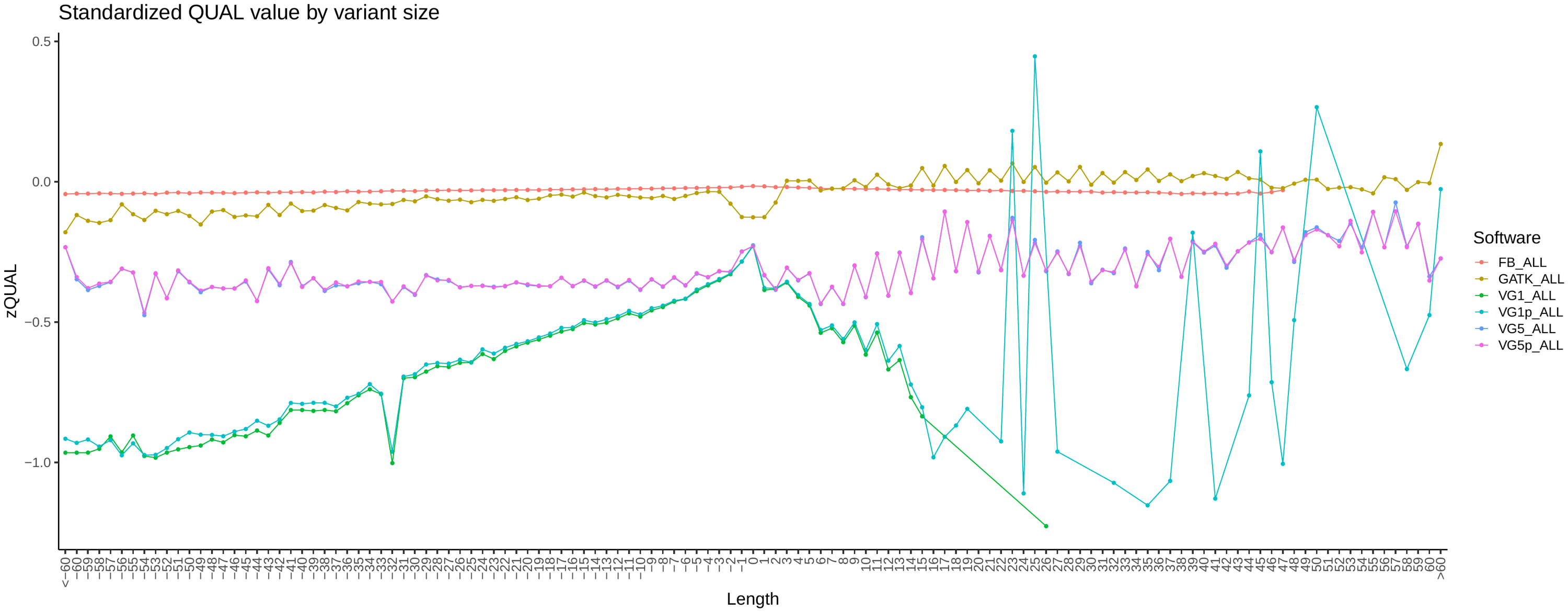

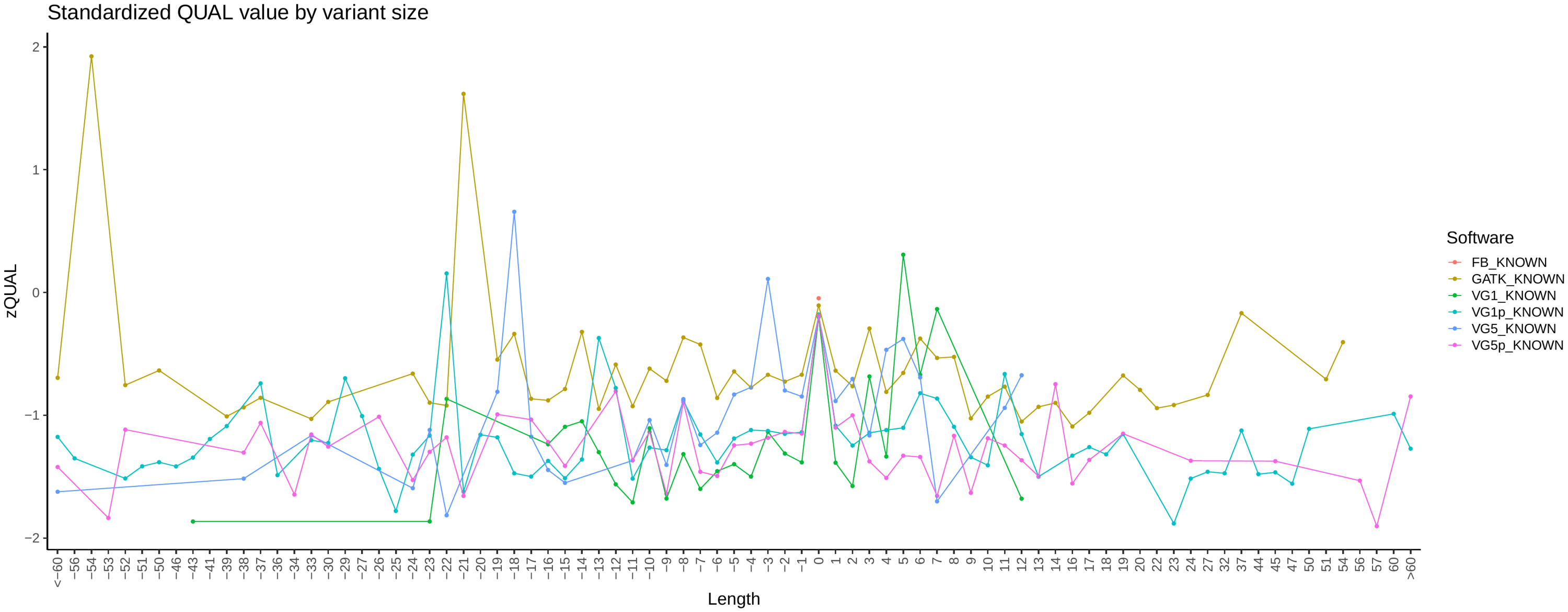

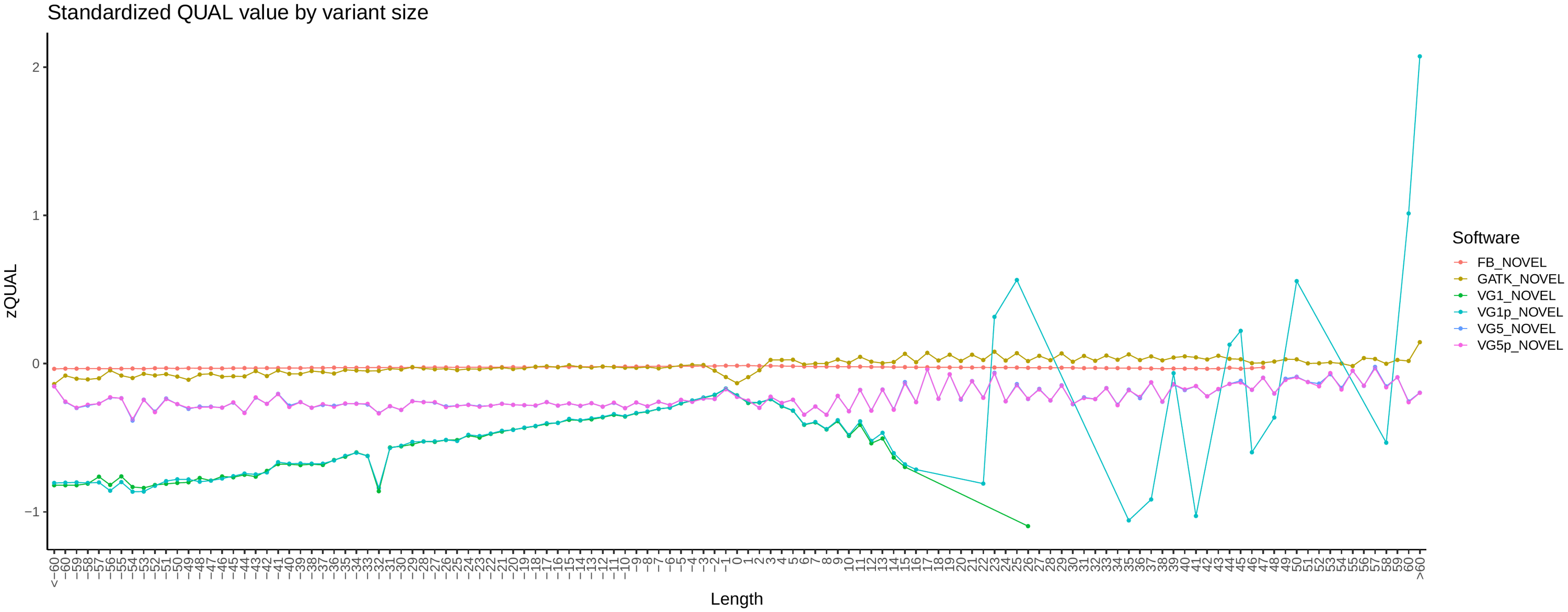

##### Variant Depth by Size

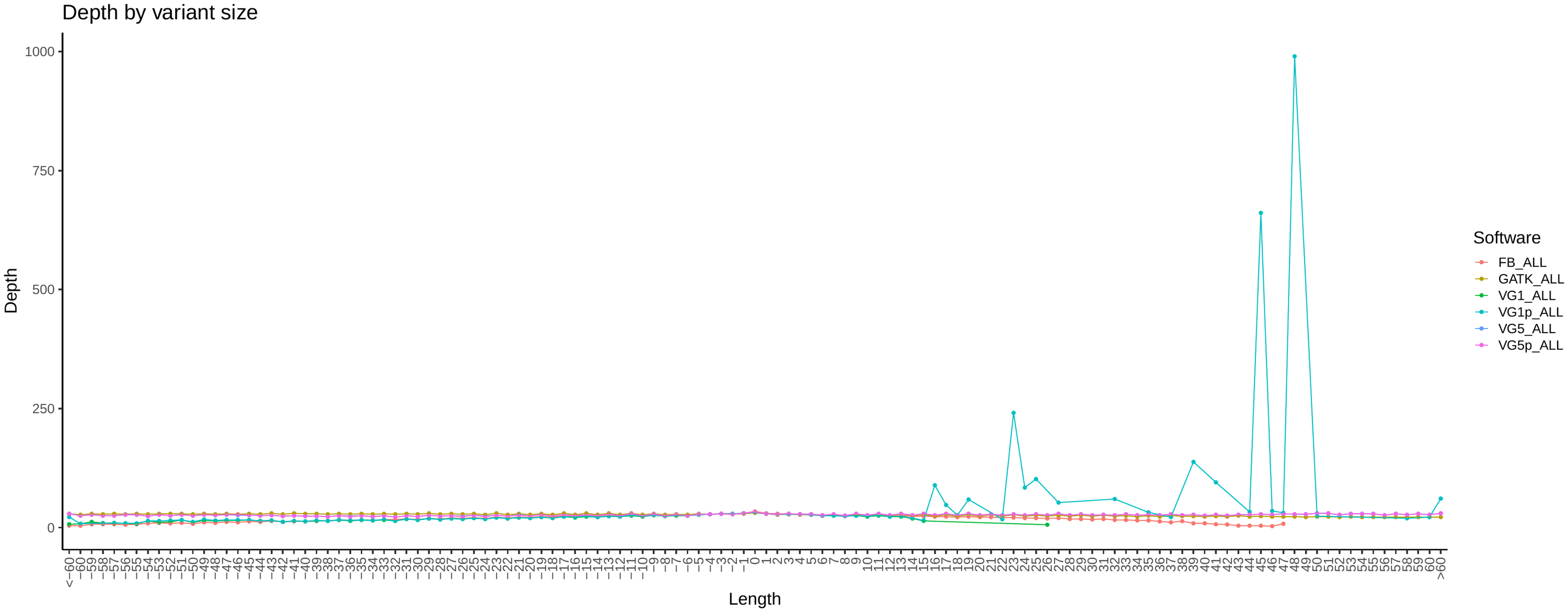

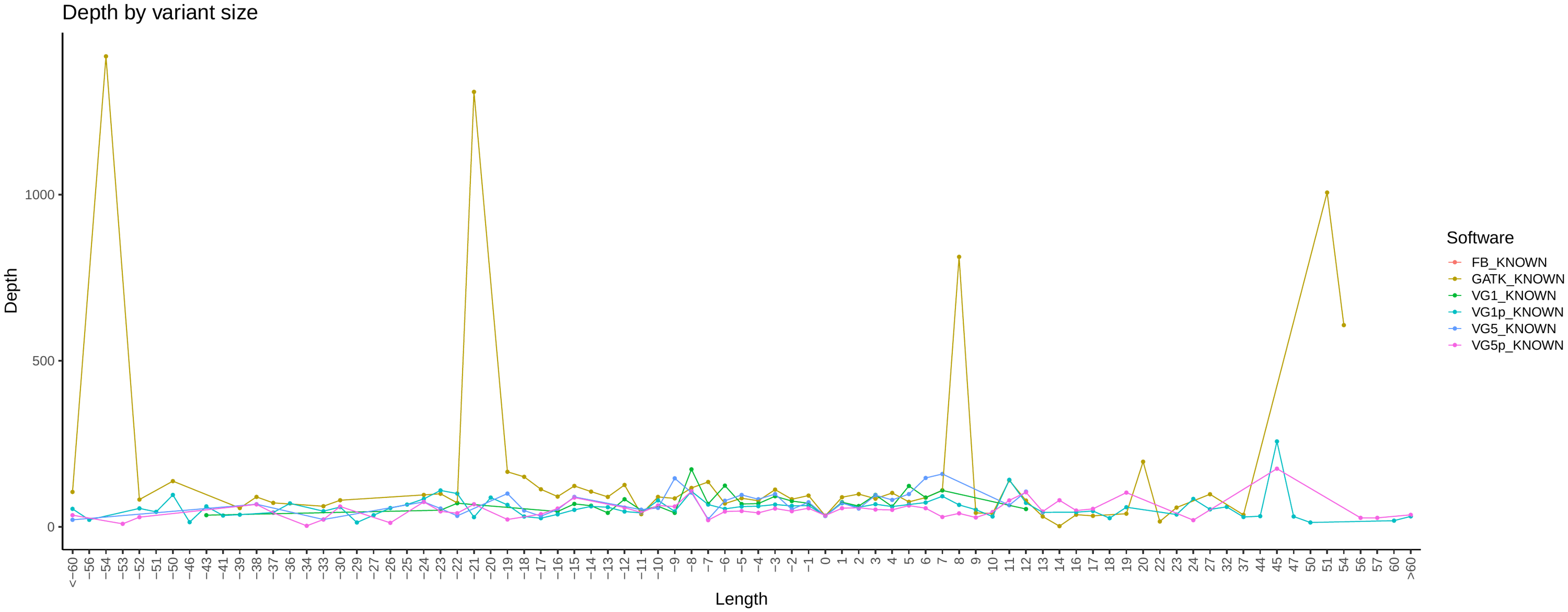

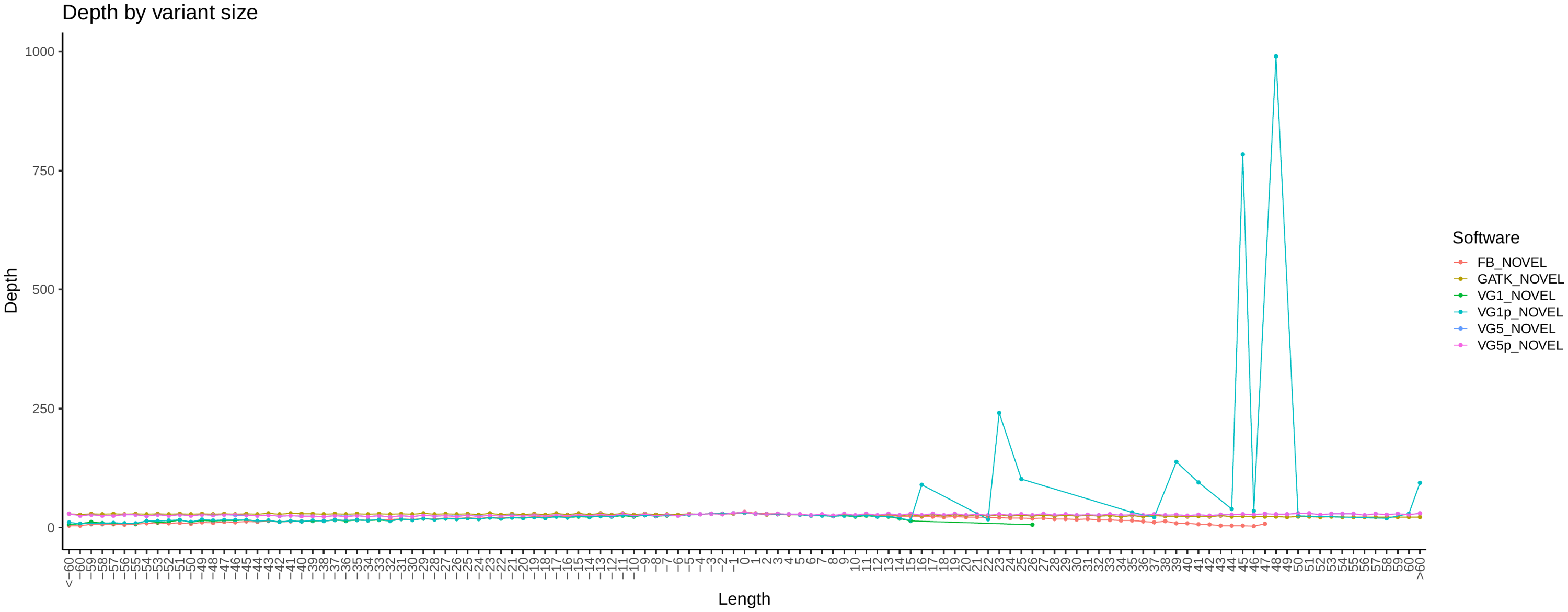

##### Variant Allelic Balance (AB)

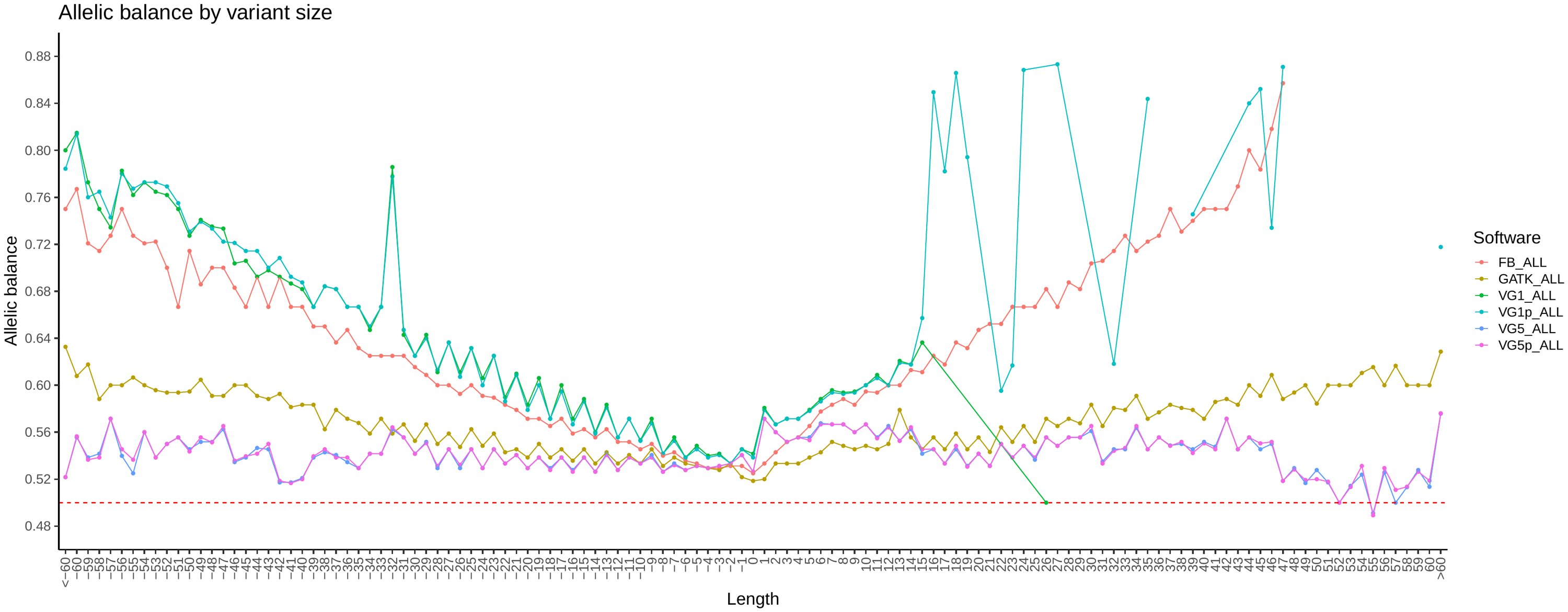

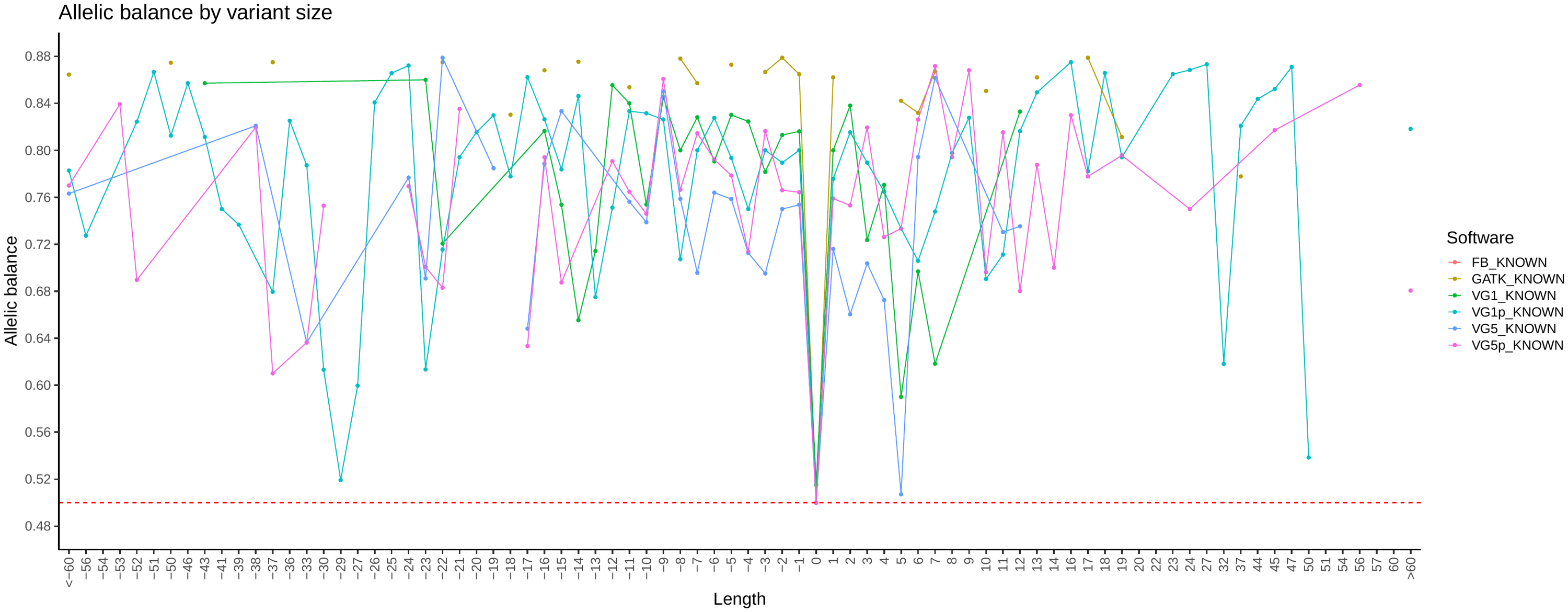

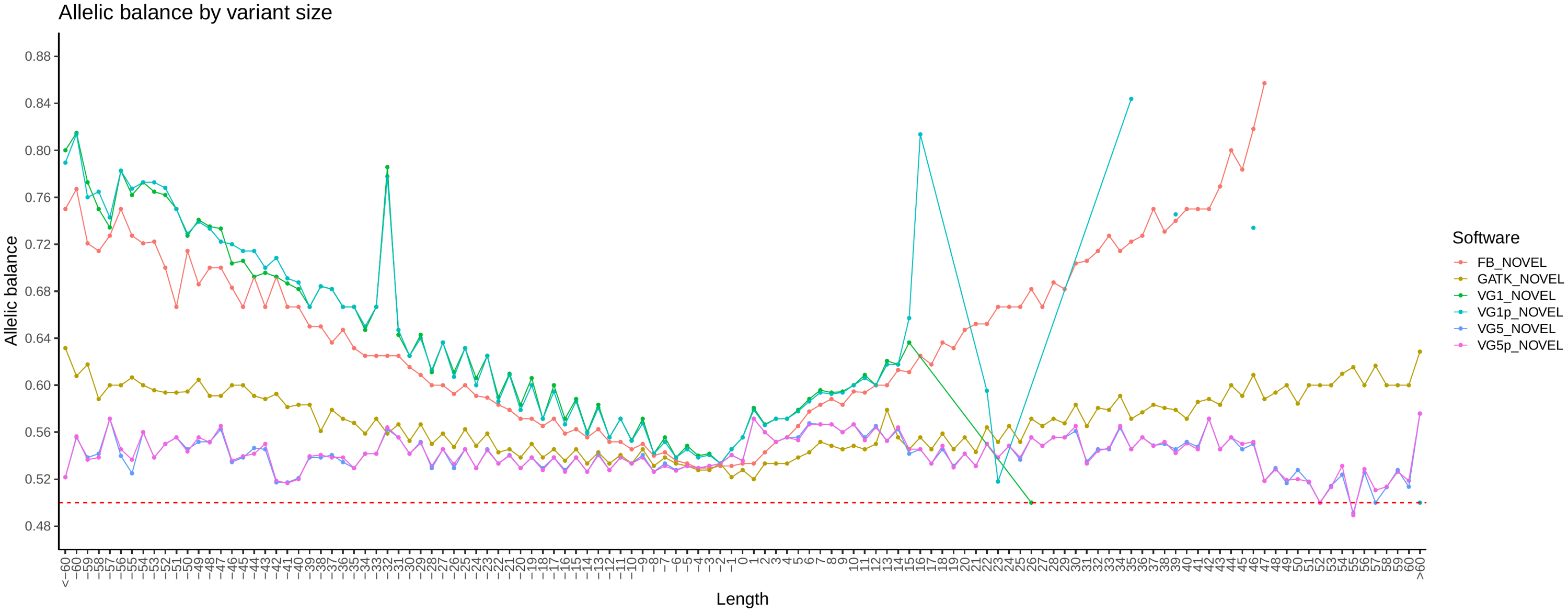

##### Variant Transition/Transversion ratio (TiTv)

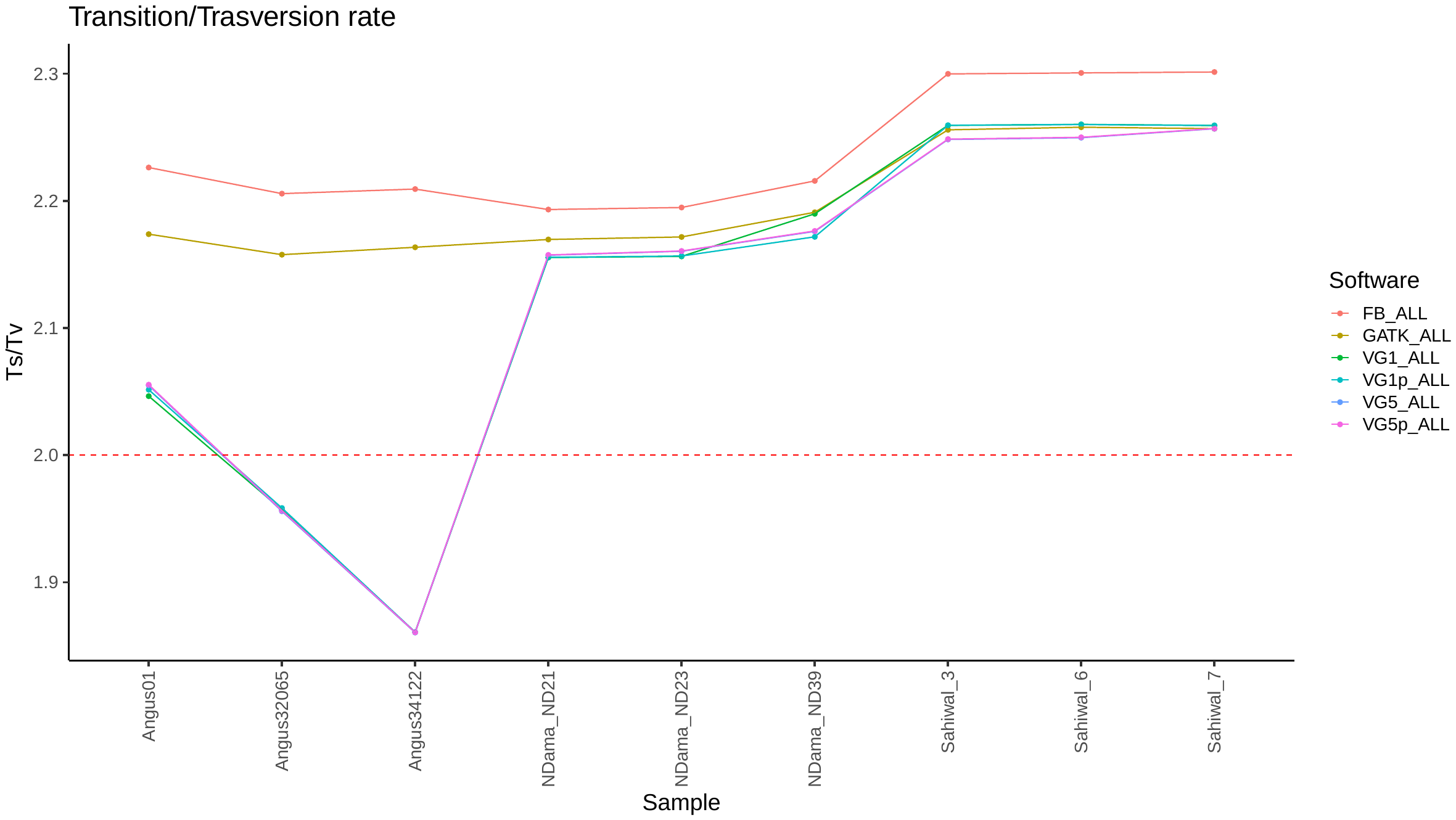

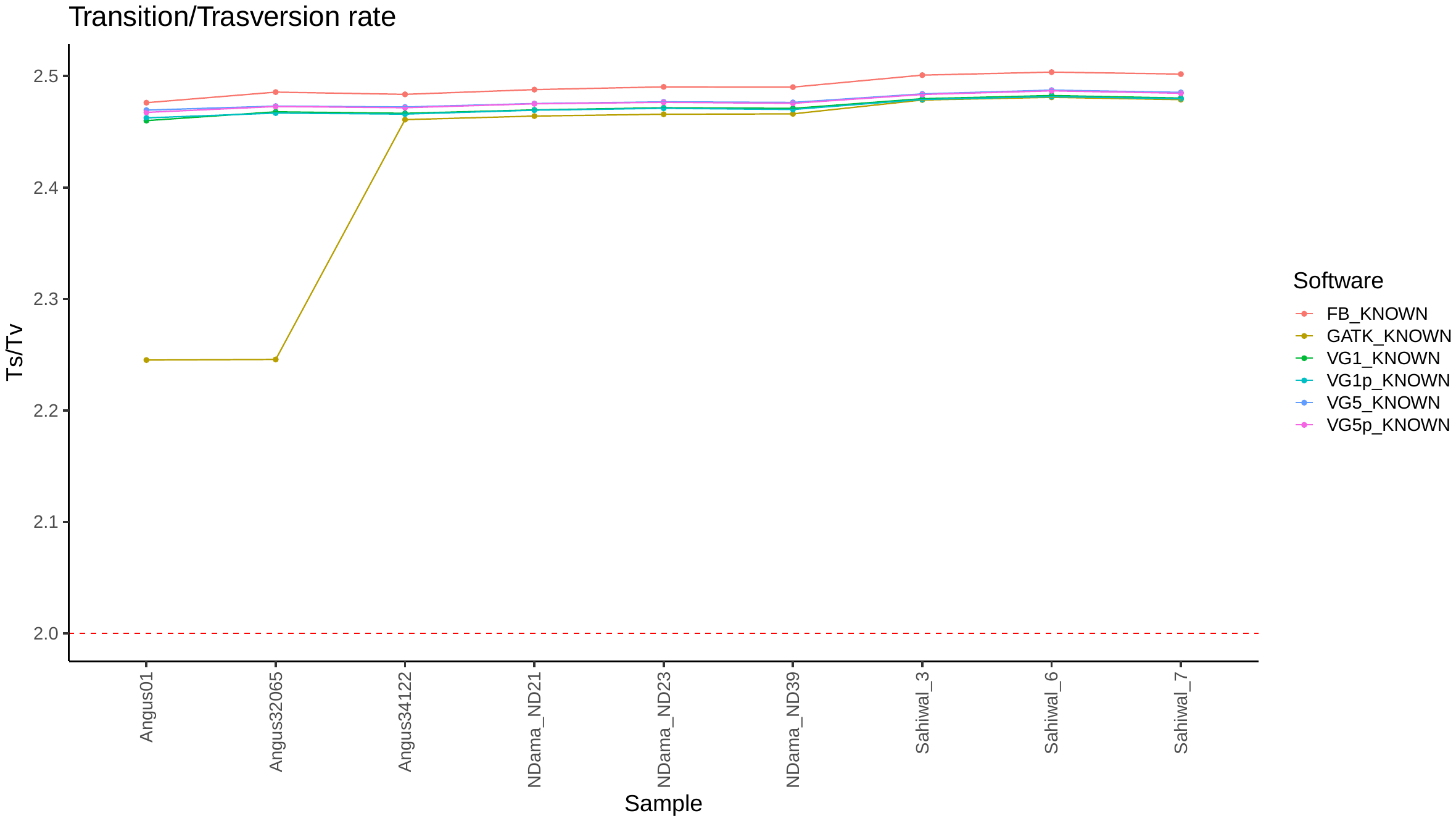

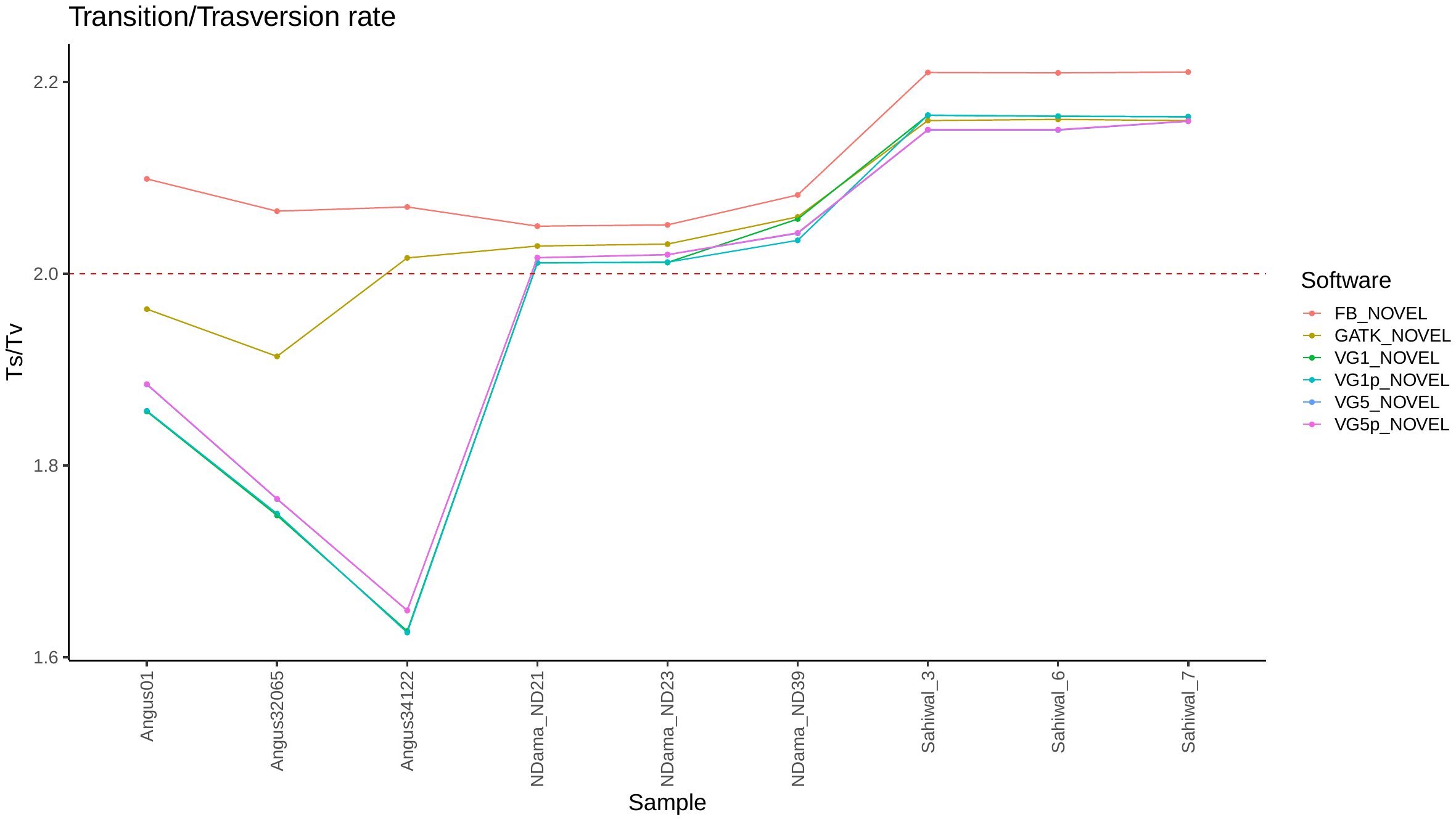

#### Scenario B: Graph-specific variants as known

##### Variant Number by size

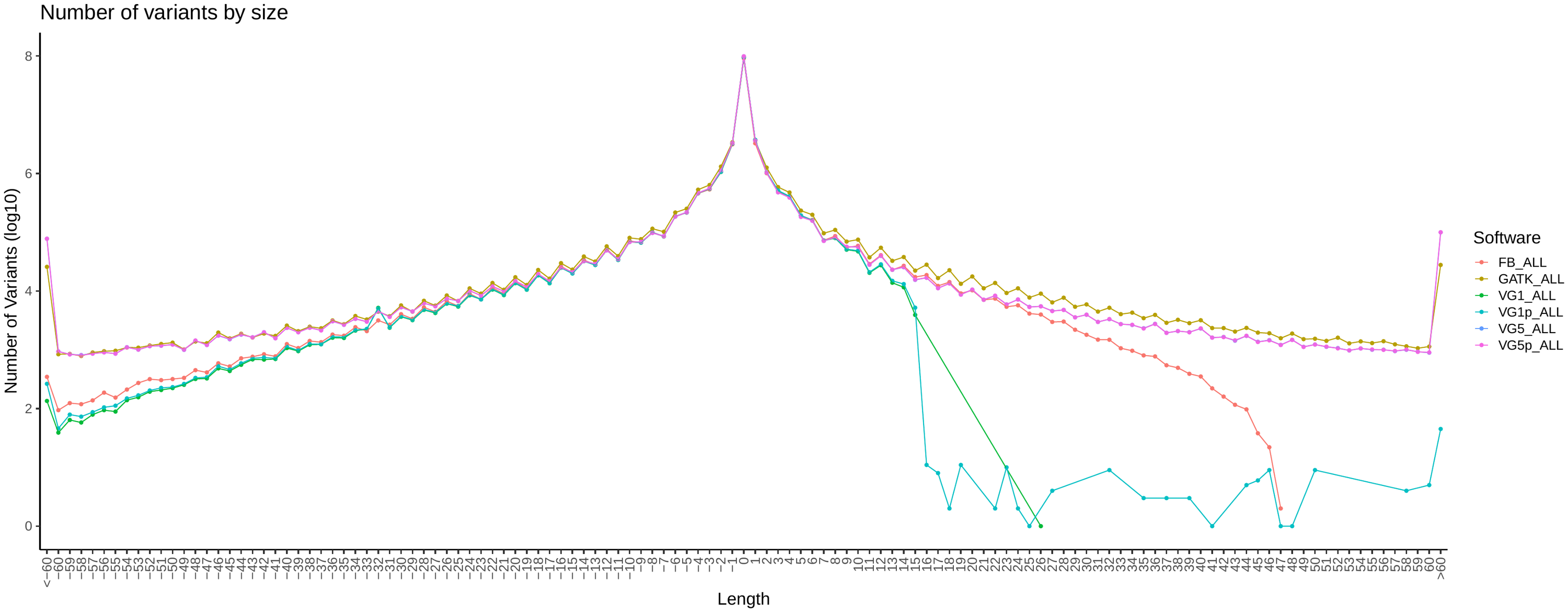

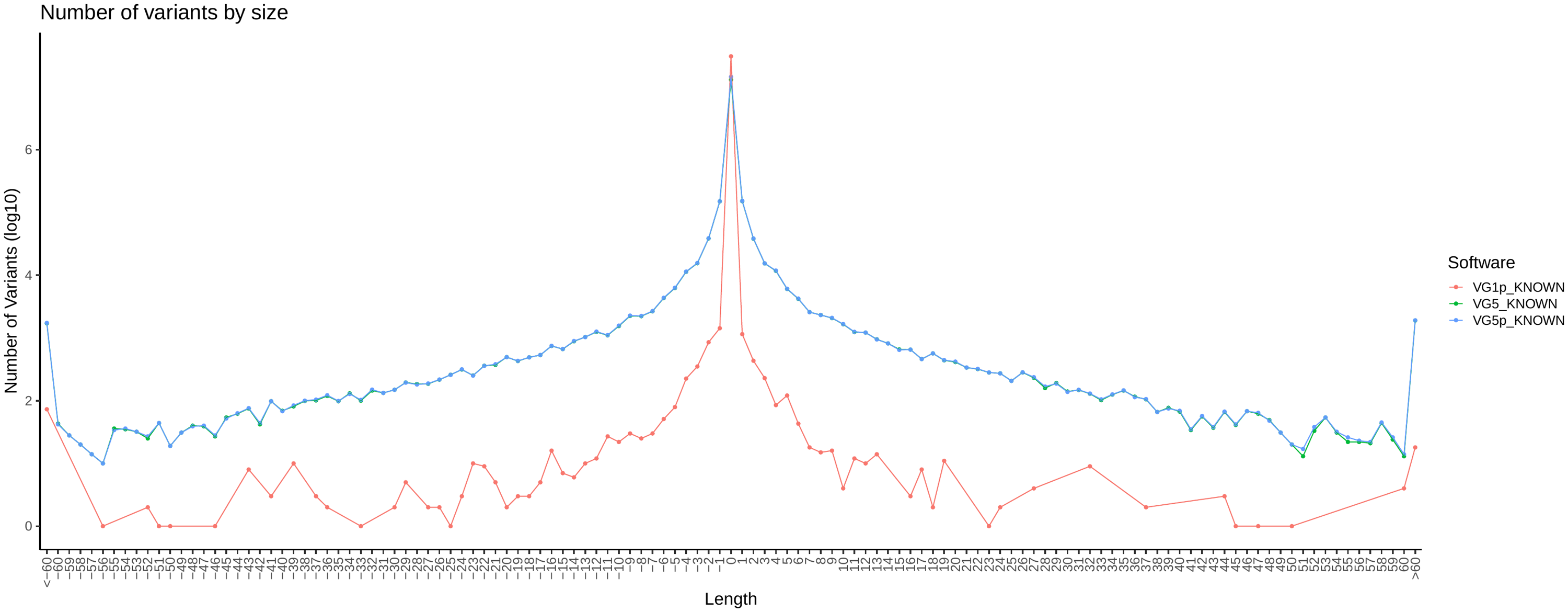

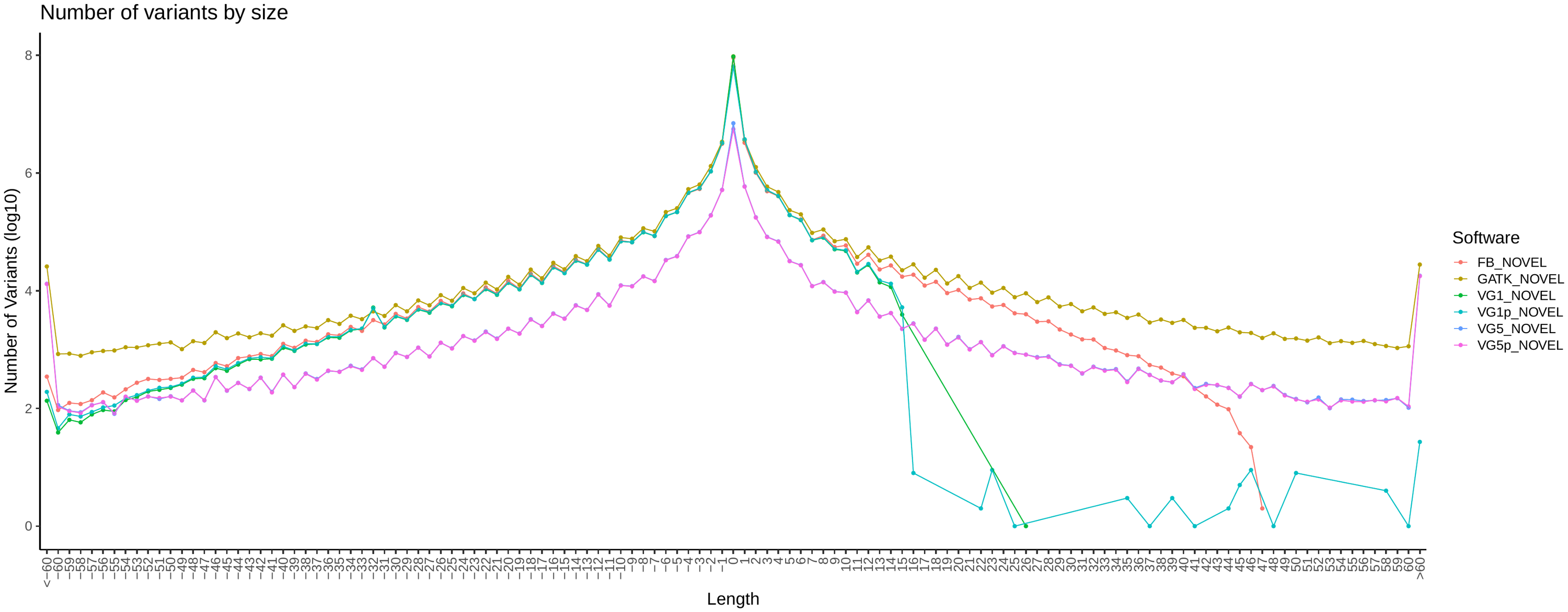

##### Variant QUAL by Size

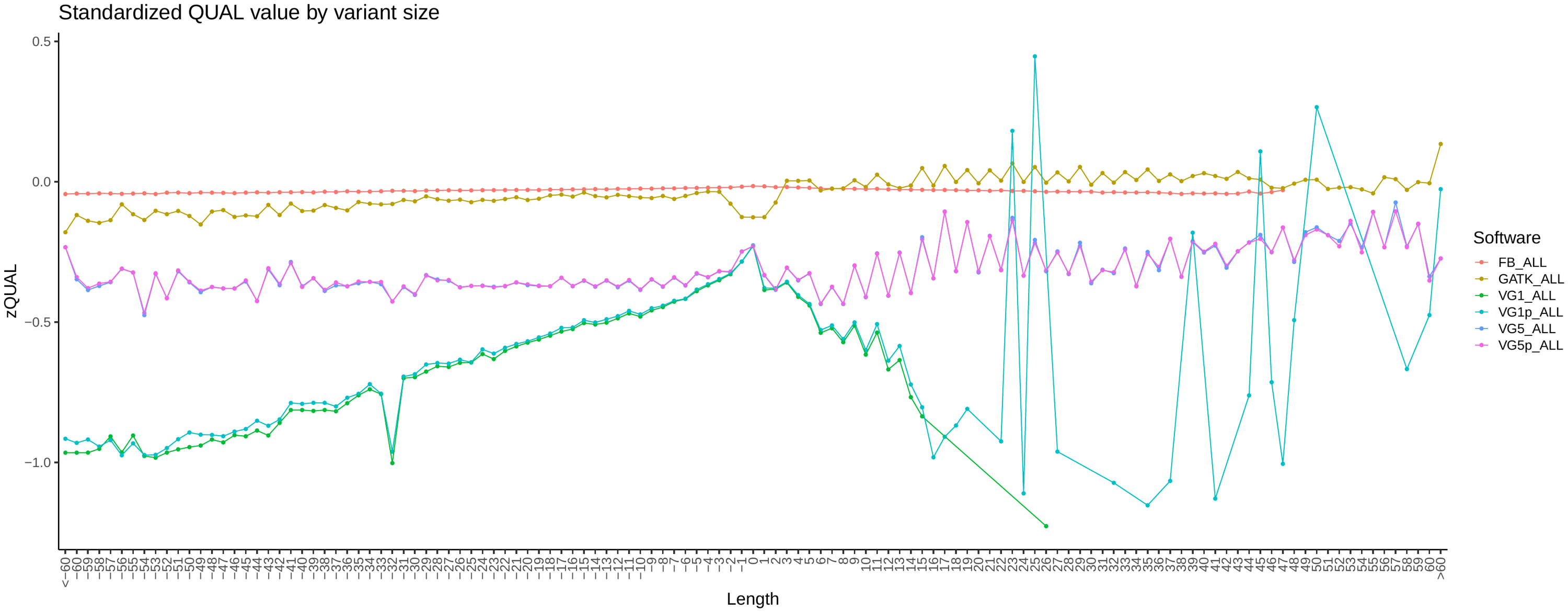

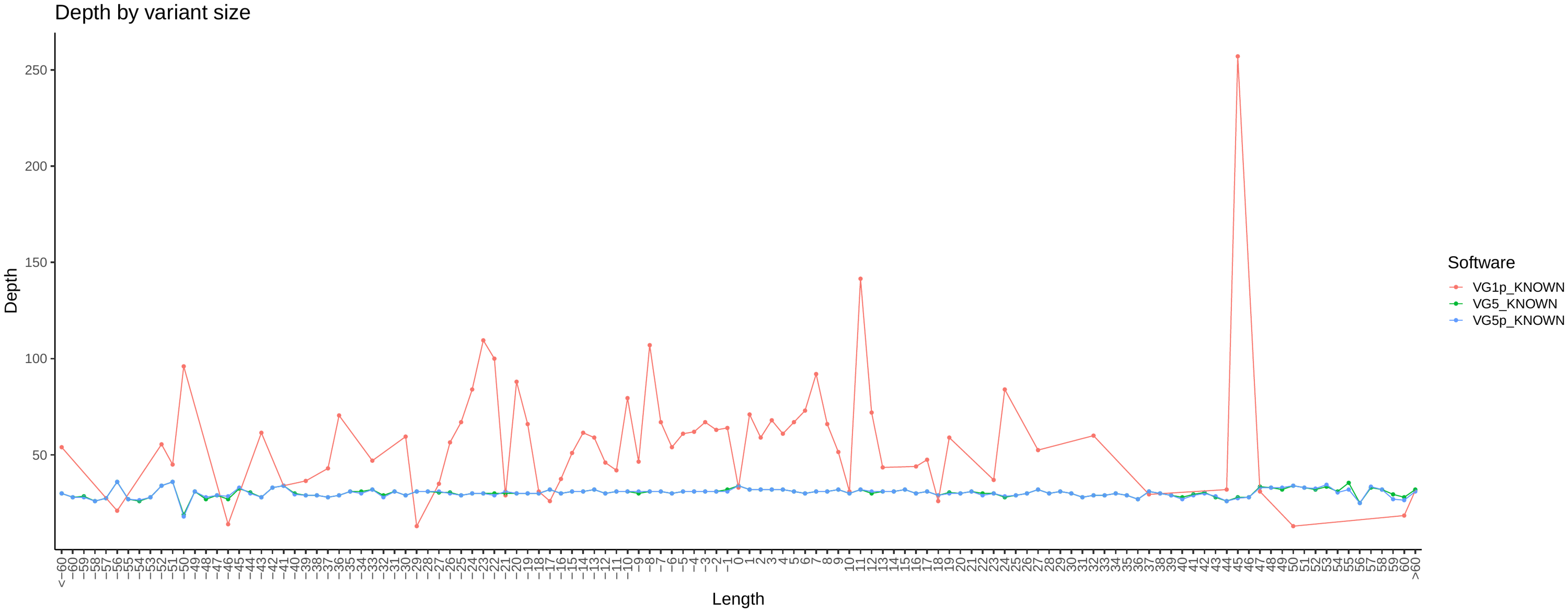

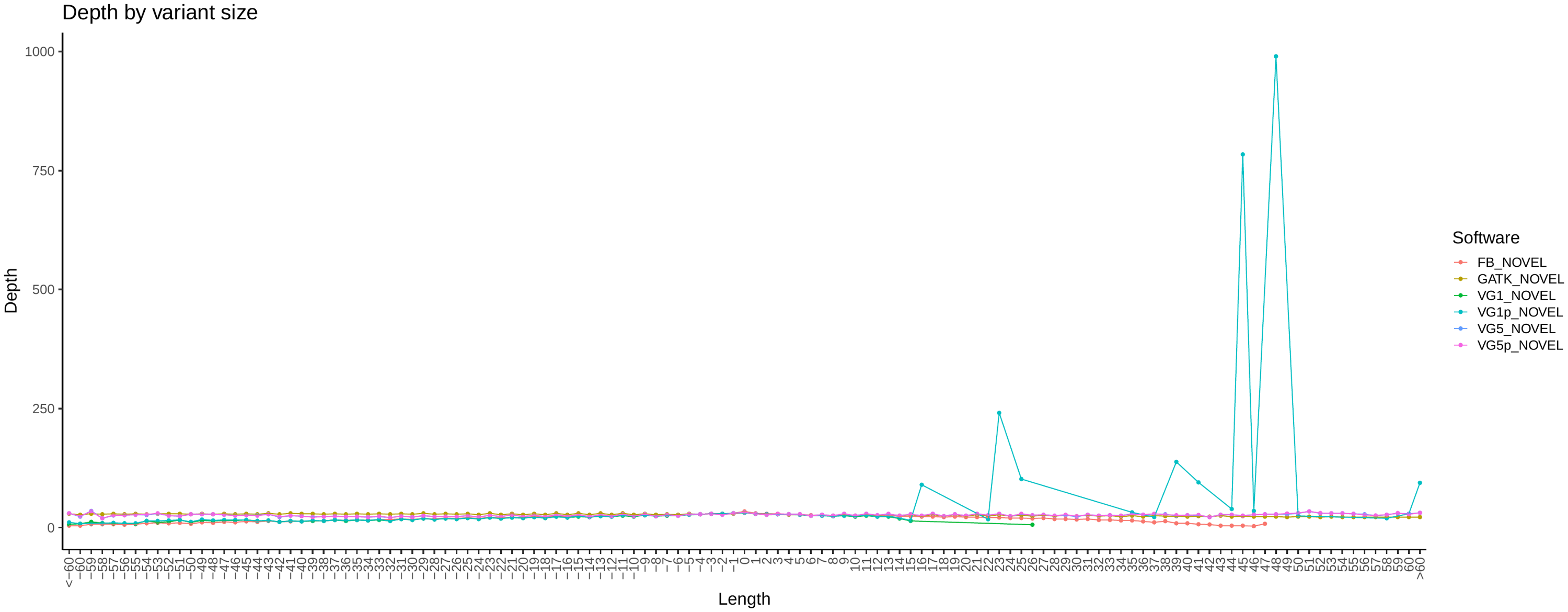

##### Variant Depth by Size

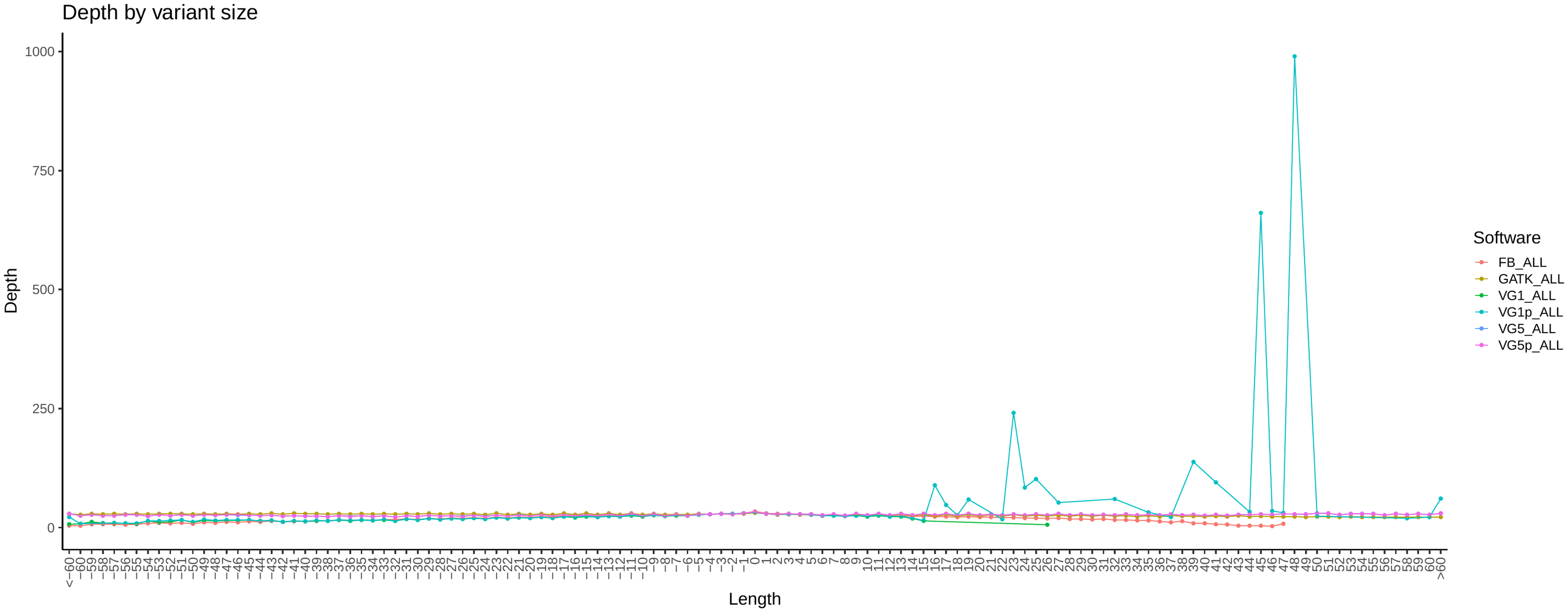

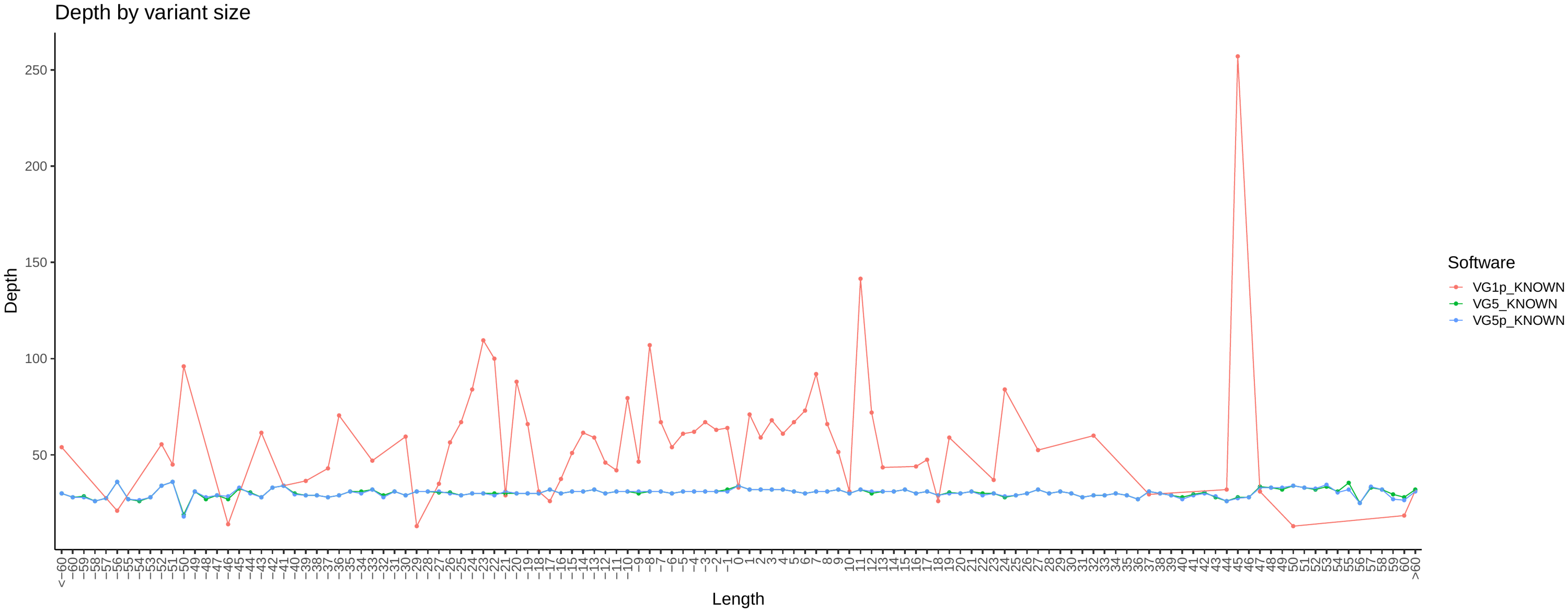

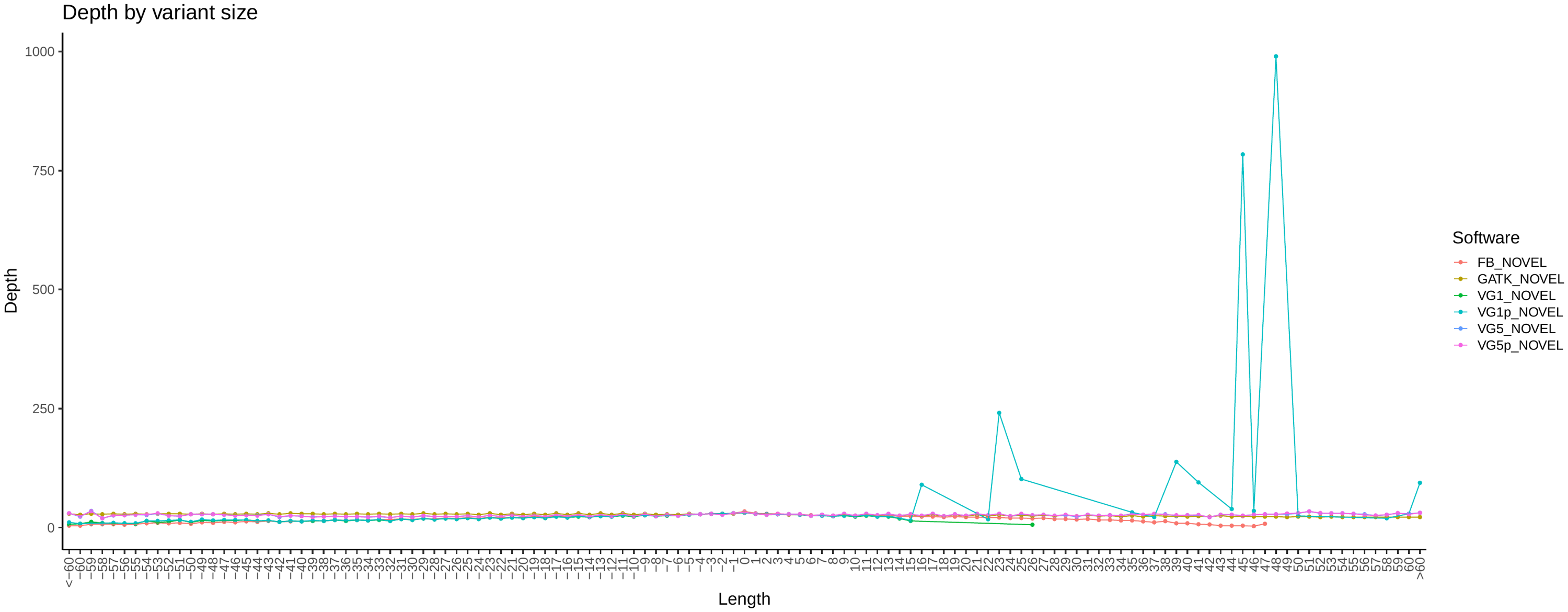

##### Variant Allelic Balance (AB) by Size

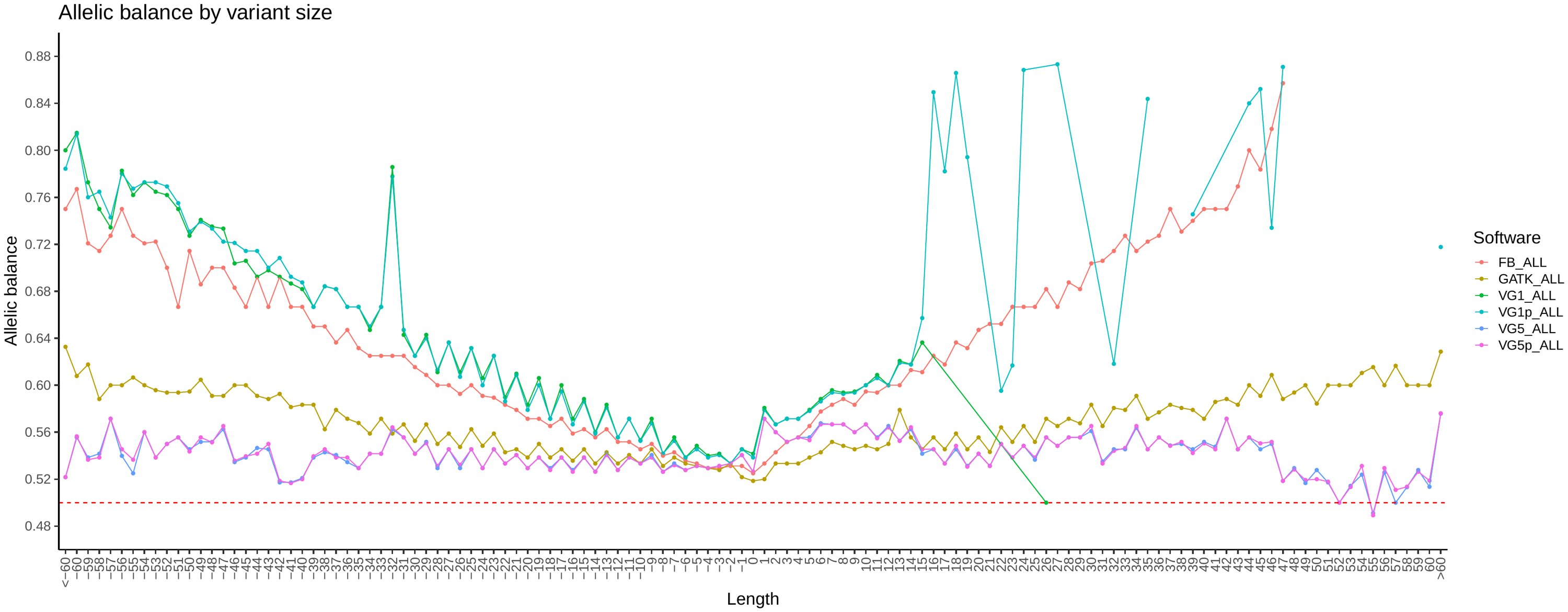

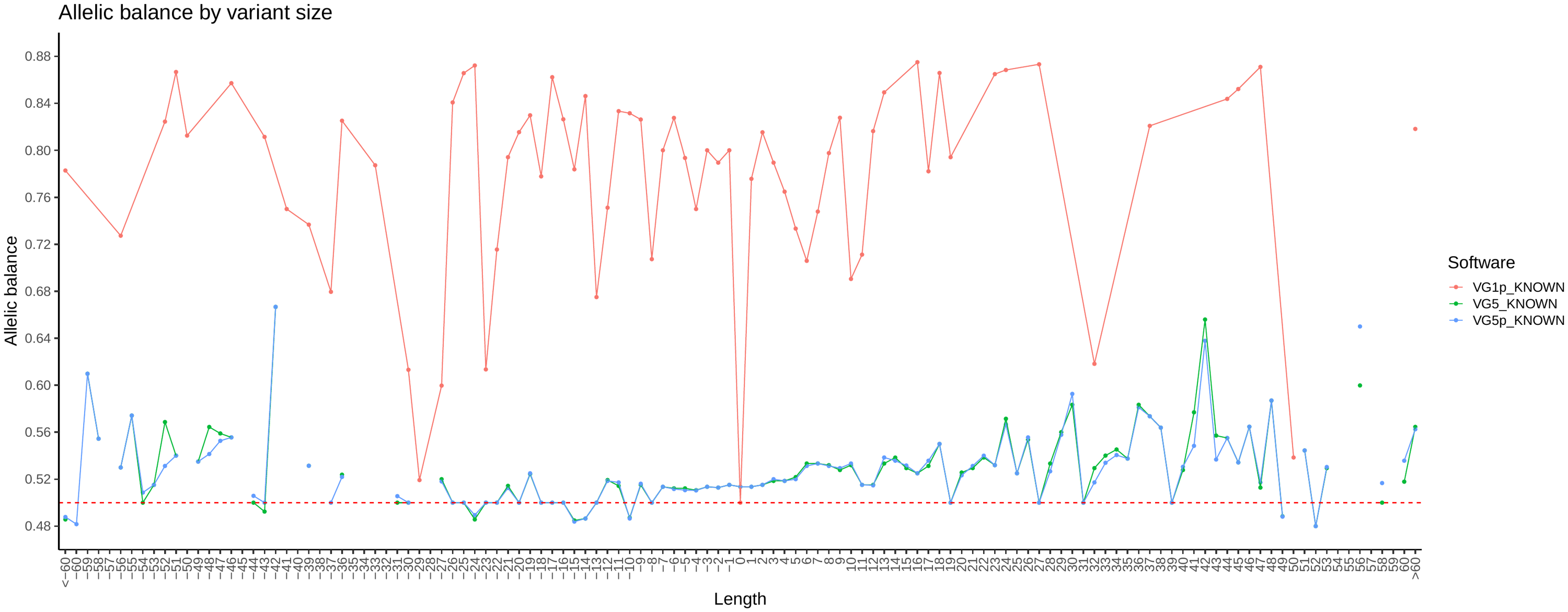

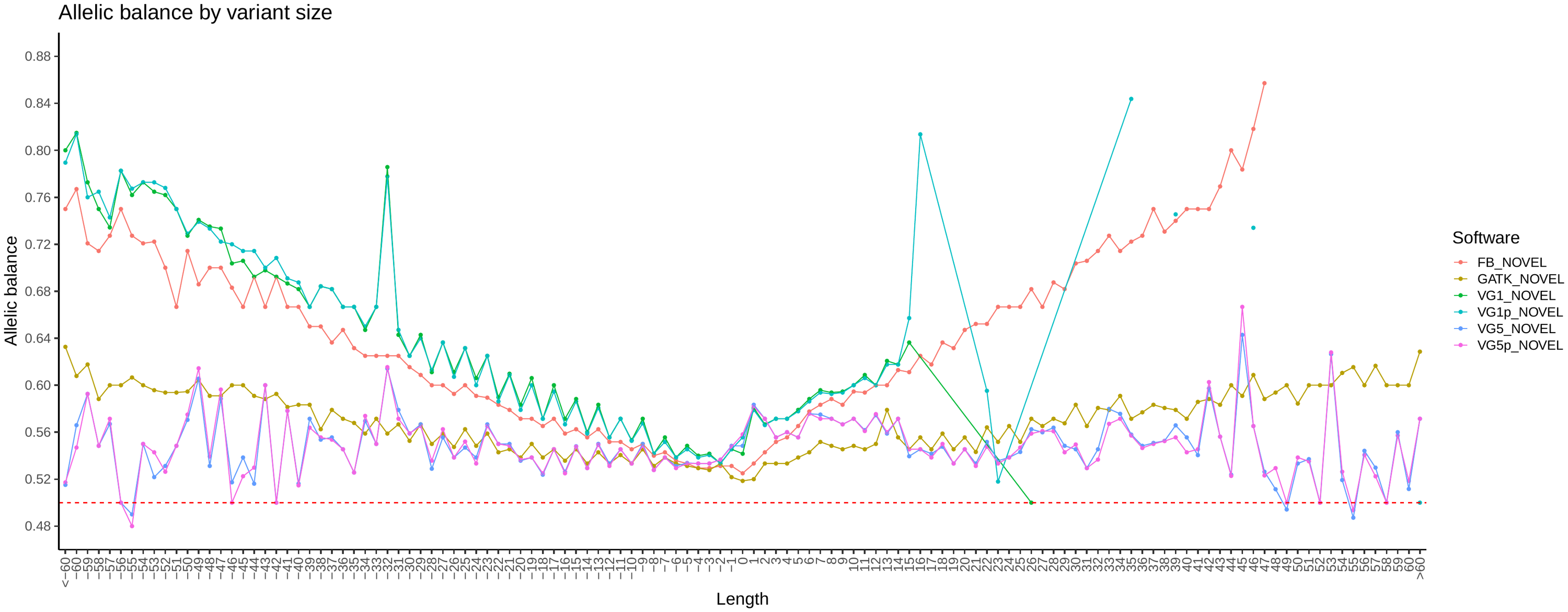

##### Variant Transition/Transversion ratio (TiTv)

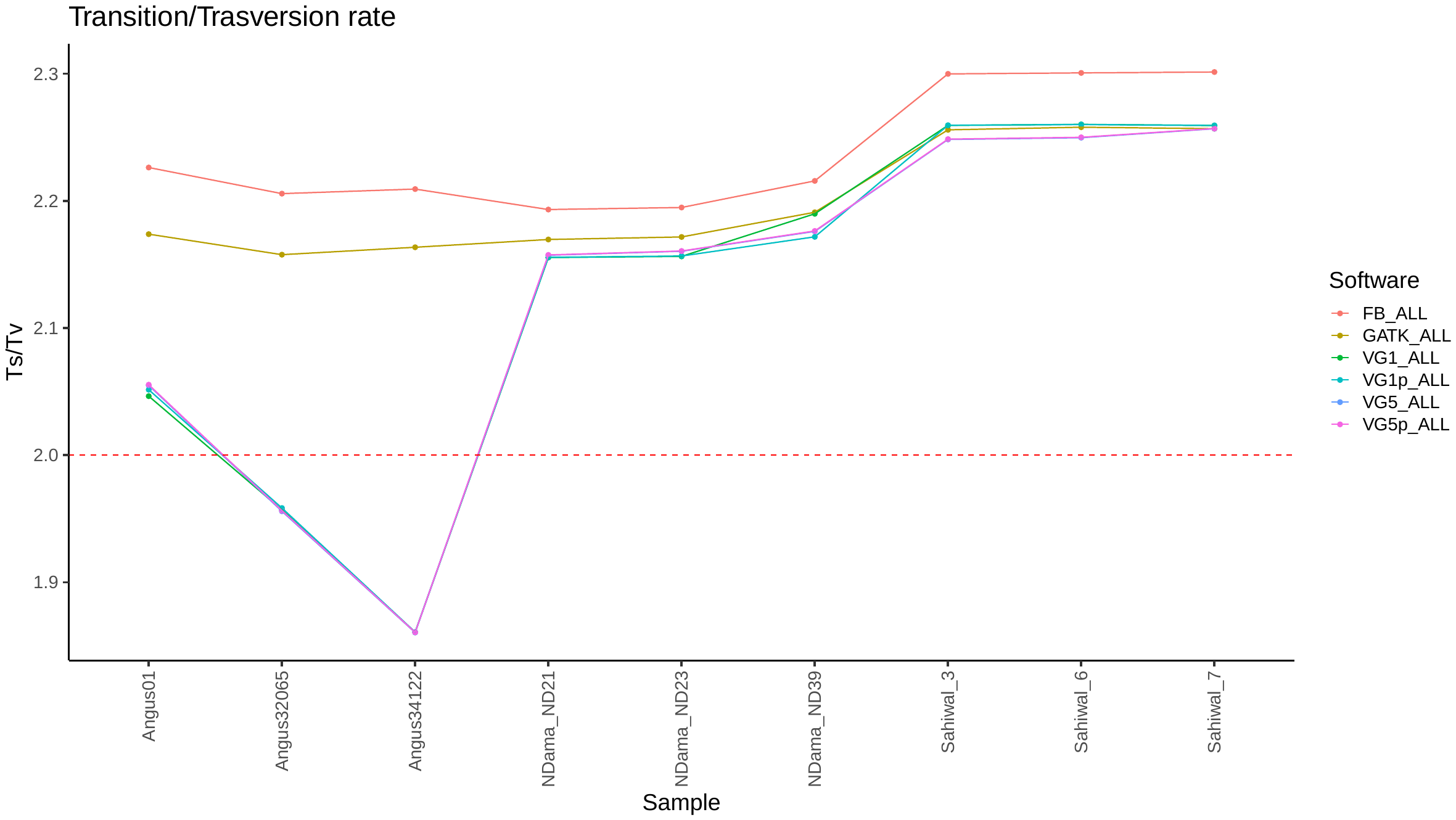

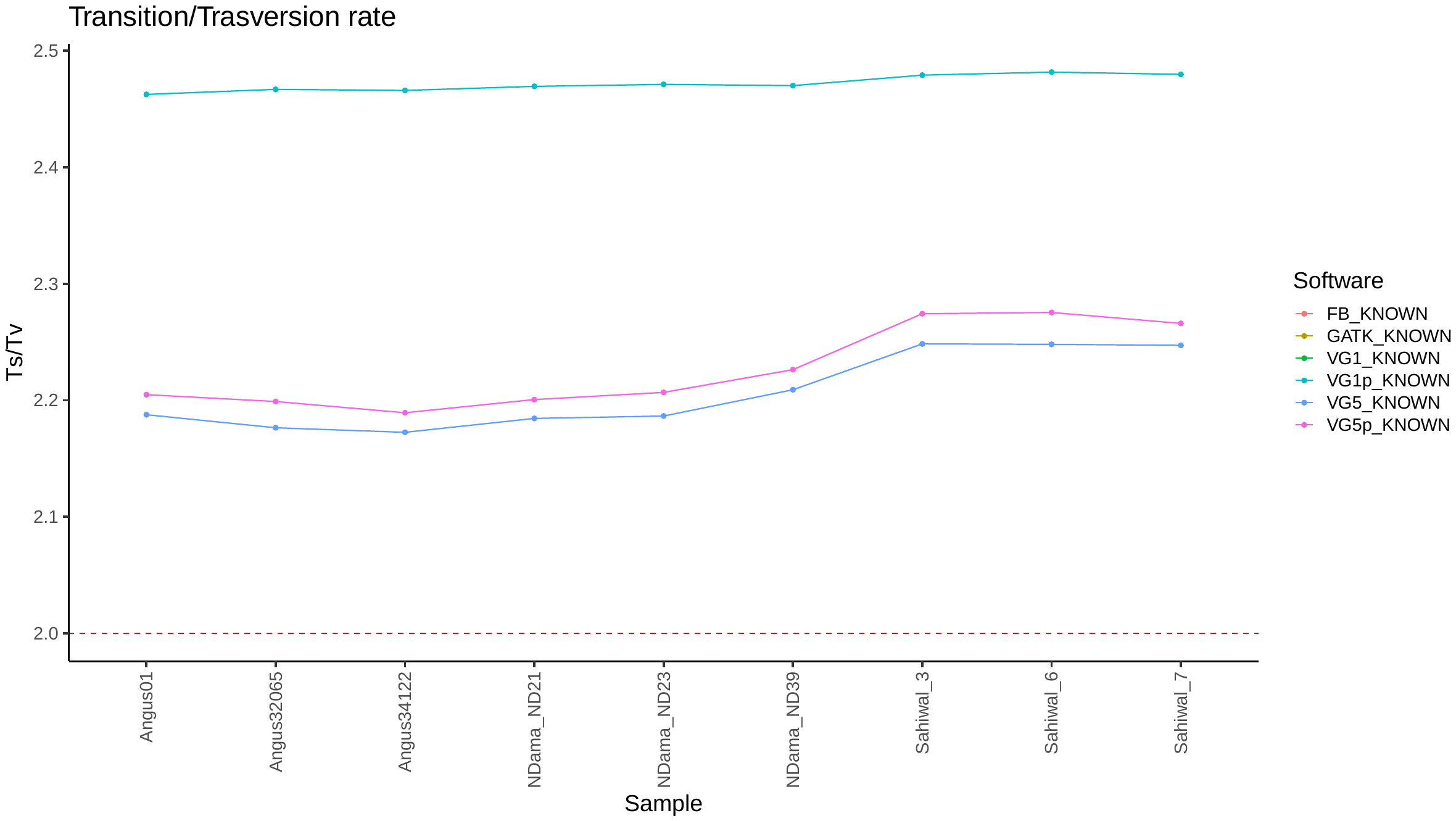

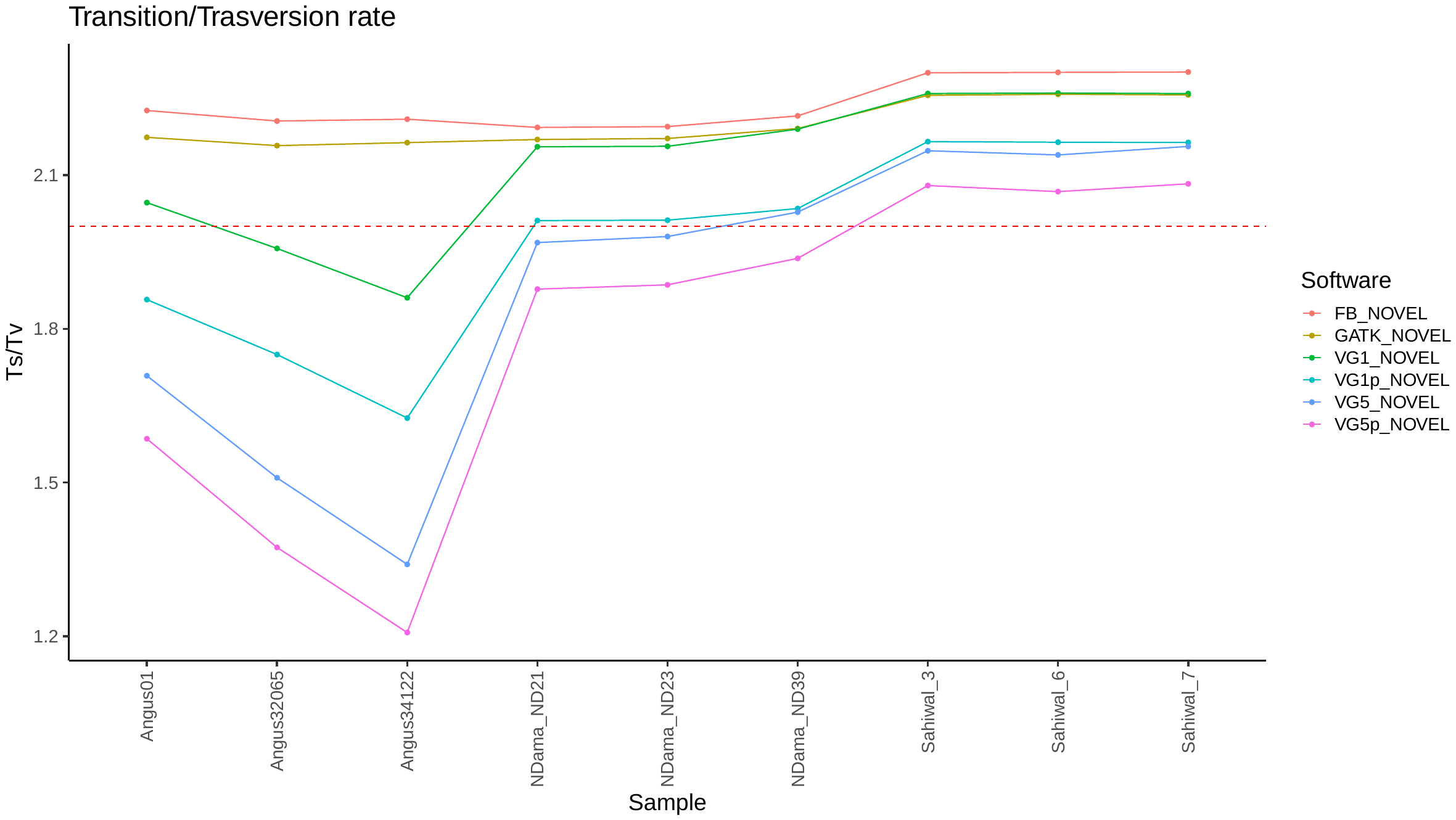
